# Extracellular matrix educates a tumor macrophage phenotype found in ovarian cancer metastasis

**DOI:** 10.1101/2022.08.11.503568

**Authors:** E. H. Puttock, E. J. Tyler, M. Manni, E. Maniati, C. Butterworth, E. Peerani, P. Hirani, V. Gauthier, Y. Liu, G. Maniscalco, V. Rajeeve, P. Cutillas, C. Trevisan, M. Pozzobon, M. Lockley, J. Rastrick, H. Läubli, A. White, O. M. T. Pearce

**Author notes:** Funding statement: E.H.P and O.M.T.P is a recipient of a Centre for Inflammation and Therapeutic Innovation (CiTI) funded studentship. E.J.T & O.M.T.P is a recipient of a CRUK & Credit Suisse career establishment award (grant code: A27947). MP is recipient of a grant of Foundation Institute of Pediatric Research Città della Speranza (grant number 27/01). P.H & O.M.T.P is a recipient of a *Barts Charity* & *Against Breast Cancer* studentship (MGU0499). Shared first authorship (E. H. Puttock, E. J. Tyler). Corresponding and senior author.

## Abstract

Recent studies have shown the tumor extracellular matrix (ECM) associates with immunosuppression, and that targeting the ECM can improve immune infiltration and immunotherapy response. A question that remains is whether the ECM is directly educating the immune phenotypes seen in cancer. We identified a tumor-associated macrophage (TAM) population correlated with poor prognosis, interruption of the cancer immunity cycle, and tumor ECM composition. To investigate whether ECM was capable of generating the TAM phenotype seen, we developed a decellularized tissue model that retains the native ECM architecture and composition. Macrophages cultured on decellularized ovarian metastasis shared transcriptional profiles with the TAMs found in human tissues. ECM educated macrophages have a tissue remodeling and immunoregulatory phenotype, inducing altered T cell function. We conclude that the tumor ECM is directly educating this macrophage population found in cancer tissues. Therefore, current and emerging cancer therapies that target the tumor ECM may be tailored to improve macrophage phenotype and their downstream regulation of immunity.

## Introduction

The remodeling of the extracellular matrix (ECM) that accompanies invasive carcinomas is associated with the establishment of an immunosuppressive environment and poor clinical response to immunotherapy^1,2^. Preclinical models where tumor ECM is targeted either through non-specific^3,4^ or specific^5^ inhibition have been shown to support anti-tumor immunity and improve response to immunotherapy. For example, broad spectrum inhibitors of ECM deposition, such as the TGF-β inhibitor galunisertib, in a preclinical model of colorectal carcinoma reduced tumor volume through improved CD8 infiltration, which in combination with anti-PDL1 improved overall survival and risk of metastasis^4^. Similarly, depleting hyaluronan, a large polysaccharide component of the tumor ECM, can improve therapy response and reduce tumor burden in preclinical studies^6–8^. These studies indicate that the tumor ECM may have a direct role in inhibiting immunity, which could occur through biophysical and/or biochemical cues. Whether the tumor ECM can directly educate the phenotype of infiltrating immune cells is not well understood but could prove important to our understanding of tumor immunity and improve immunotherapy efficacy.

In our previous work, we identified a specific tumor ECM signature that associated with disease progression and immunosuppression, identified in high-grade serous ovarian cancer (HGSOC)^1^, and shared by 12 other solid carcinomas, including those with limited treatment options such as pancreatic, lung, and esophageal cancers. This ECM signature associated with immunosuppressive cell phenotypes, but did not positively correlate with cytotoxic immune cell phenotypes, and negatively correlated with CD8 T cells^9^. Building on these studies, we applied computational algorithms CIBERSORTx and xCell to these previously gathered datasets^1^ to further look at how specific ECM molecules associate with immune cell infiltrate. This analysis identified a population of tumor-associated macrophages (TAMs) associating with five ECM molecules that are predictive of poor prognosis, and have been identified previously in signatures that associate with immunosuppression, poor prognosis, and failure of immunotherapy response^10^. These macrophages were characterized by the gene signature for M0 macrophages that are used in CIBERSORTx and xCell. Originally the signature for M0 macrophages was used to define a non-activated macrophage phenotype, however more recent studies indicate that the signature is also a feature of a poorly characterized macrophage population that associates with poor prognosis in hepatocellular^11^, sarcoma^12^, breast^13,14^, and lung^15^ cancers. TAMs likely exist as a spectrum of subtypes influenced by their interaction with malignant and immune cells and secreted factors during infiltration into the tumor tissue^16,17^. At the poles of TAM polarization, classically activated or M1-like macrophages support inflammation and associate with better outcomes, whereas alternatively activated, or M2-like macrophages suppress anti-tumor immunity. These macrophage phenotypes can be generated through cytokines released within the local microenvironment, for example macrophages that exhibit more M1-like phenotype are stimulated through interferon-gamma (IFN-γ), whereas macrophages that exhibit more M2-like phenotype are stimulated through interleukins (IL) such as IL-4 and IL-13. Whether the tumor ECM is also a factor in the generation of TAM populations is not well understood. To answer this question, we made a decellularized tissue model of omental metastasis from whole patient samples, which comprises a heterogeneous ECM derived from a variety of tumor microenvironment (TME) cell types, where we found monocytes differentiate into TAMs that overlap with the M0 and M2 signatures. These ECM derived TAMs have gene programs associated with T cell activation and tissue remodeling. This study demonstrates that tumor ECM can directly educate TAMs found in ovarian cancer tissues that associate with poor prognosis and suggests that strategies that target ECM may also alter immune cell phenotype as well as their infiltration within the tumor microenvironment.

## Results

### An underexplored HGSOC macrophage subtype associates with a prognostic tumor ECM signature

Several bioinformatics studies connect immune cell infiltration with the composition of the tumor ECM^1,2,18^. To investigate changes that occur to immune cell phenotypes in ovarian metastasis we used computational algorithms CIBERSORT, CIBERSORTx and xCell which are validated to predict and deconvolute immune cell abundances in tumor tissue from patient biopsies (Figure 1A). To investigate ECM molecules differentially expressed and associated with immune cells we used gene set enrichment analysis and differential gene expression in an omental metastasis dataset testing their accuracy at estimating immune cell populations in a published dataset of omental metastasis of HGSOC samples where transcriptomic, proteomic, and immune cell count data was available^1^ (Figure 1B-E, Supplemental Figure 1, and Supplemental Methods). As anticipated, deconvolution of our bulk RNA-sequencing (RNAseq) dataset using analytical tools CIBERSORTx and xCell (which were designed to analyze RNAseq data) accurately estimated the abundance of CD8^+^ (R = 0.57 and 0.60, respectively), CD68^+^ (R = 0.36 and 0.46, respectively), CD3^+^, CD4^+^, and CD45RO^+^ immune cell counts. CIBERSORT, an earlier version designed to analyze microarray data, did not accurately estimate abundances and was not used further. We next compared the immune signatures generated by both CIBERSORTx and xCell against a previously reported tumor ECM signature and measure of tumor desmoplasia^1^ in the same cohort of patients identifying several correlating immune cell types including macrophages, B cells, several T cell subsets, dendritic cells, basophils, and neutrophils, which together confirmed our previous results using a less comprehensive set of immune cell signatures (Supplemental Figure 2A-B). Of note, macrophages correlated highly with these signatures, and the majority of intra-tumoral macrophages are thought to exhibit an M2-like phenotype which correlates with poor prognosis in several malignancies, including ovarian cancer^19,20^. We identified M2 macrophages as a predominant signature present in all our HGSOC patient samples (Supplemental Figure 1A-C). However, we did not find a significant association for alternatively activated ‘M2-like’ macrophages with the level of disease (Supplemental Figure 2A-B), an observation which has been reported previously^21,22^, which may result from heterogeneity between patient samples. There was a strong correlation with the CIBERSORTx ‘M0 macrophage’ signature and ‘Macrophage’ xCell signature, both of which are derived from the Immune Response In Silico (IRIS) gene expression dataset for macrophages after 7 days of differentiation from monocytes^23–25^. The CIBERSORTx M0 macrophage gene signature has been more recently associated with tumor promotion and poor prognosis in multiple cancers^11–15^ but it is unclear if the M0 signature is describing a distinct population of macrophages or (more likely) multiple populations of macrophages which do not fit well with classical and non-classical macrophage signatures. The M0 population has been previously reported in murine HGSOC mouse models^26^, however, this is the first observation of M0 macrophages within the TME of patients with HGSOC.

**Figure 1.**
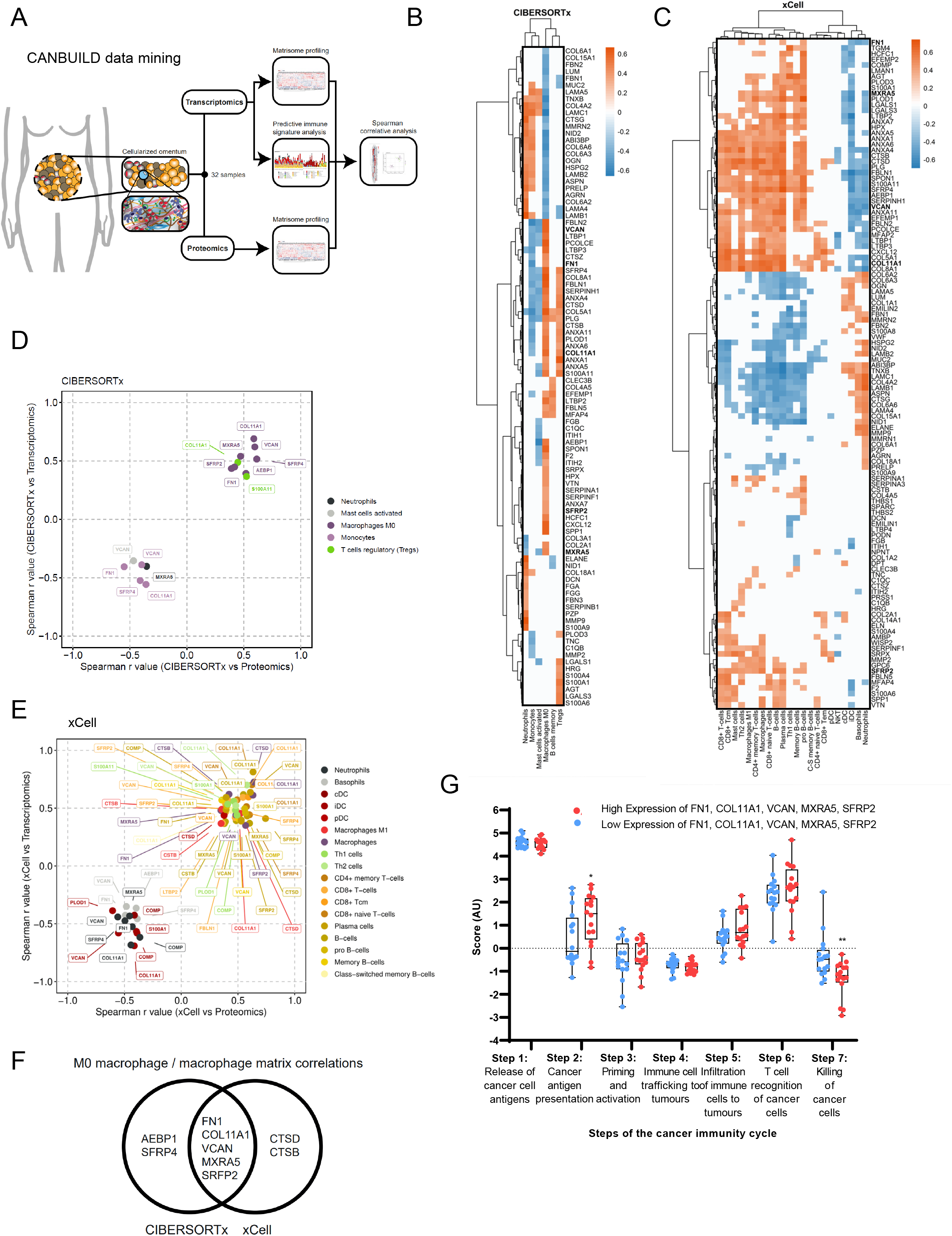
M0 macrophages are enriched in highly diseased HGSOC samples and significantly correlate with a matrisome signature. **A)** Schematic of bioinformatics pipeline. **B)** Heatmap of ECM proteins significantly associated with immune cells from CIBERSORTx analysis. Spearman’s r values representing correlation between immune abundances and protein expression levels are plotted according to the color scale, with positive correlations in red and negative in blue. White colored associations are insignificant with a correlation p > 0.05, N = 32 HGSOC samples. Ordered by unsupervised clustering. **C)** Heatmap of ECM proteins significantly associated with significant immune cells from xCell analysis. Spearman’s r values representing correlation between immune abundances and protein expression levels are plotted according to the color scale, with positive correlations in red and negative in blue. Cream colored associations are insignificant with a correlation p > 0.05, N = 32 HGSOC samples. Ordered by unsupervised clustering. **D-E)** Scatterplot of significant protein and gene correlations with associated immune cell types using **D)** CIBERSORTx and **E)** xCell in HGSOC. Spearman’s regression analysis was completed to generate plotted r values. Points are colored based on their associated immune cell type as depicted in the key. **F)** Venn diagram showing overlap of ECM molecules significantly correlated with CIBERSORTx M0 macrophage signature and xCell macrophage signature **G)** Boxplot of predicted tumor immune phenotype stage scores by TIP, with HGSOC samples separated by high (red) and low (blue) protein expression levels of FN1, VCAN, MXRA5, COL11A1, SFRP2. Scores are based upon expression of signature genes and represent activity levels. N = 32 (16 low and 16 high). Two-way ANOVA significance between each group is presented as **p < 0.01 and *p < 0.05.

Next, we identified correlations of immune cell populations with ECM molecules. To identify significant immune-ECM associations, we integrated sample matched CIBERSORTx and xCell immune signatures against gene and protein expression values for ECM molecules found in the matrisome database^27^, using Spearman correlative analysis (Figure 1B-E, Supplemental Figure 3). 16 ECM molecules were significantly positively associated with immune cell signatures at gene and protein levels, including macrophages, T cells, and B cells (Supplemental Figure 4). Five ECM molecules, fibronectin (FN1), versican (VCAN), matrix remodelling associated 5 (MXRA5), collagen 11 (COL11A1), and secreted fizzled related protein 2 (SFRP2) were consistent between CIBERSORTx and xCell and correlated with CIBERSORTx M0 Macrophage and xCell Macrophage signatures (Figure 1F, Supplemental Table 1). Individually these five molecules associated with several immune cell types (Figure 1D-E), but only the association with M0 macrophages was shared by all five molecules across both the CIBERSORTx and xCell approaches (Figure 1F). To explore the association of these five M0 macrophage-associated ECM molecules with the immune microenvironment in cancer, we tested whether the signature correlated with the cancer immunity cycle (Figure 1G), a series of seven stepwise events that help inform how an immune response can be activated against an established tumor^28^. To do this, we utilized TIP^29^ (Tracking Tumor Immunophenotype), which showed a significant reduction in cancer killing activity (step 7 of the cancer immunity cycle, Figure 1G) suggesting that this ECM signature may be related to the presence of immunosuppressive phenotypes (Figure 1G). TIP analysis also showed an increase in step 2, cancer antigen presentation, but no further significant differences between the other five steps of the cycle (Figure 1G). Laser capture microscopy of tumor and stroma areas from two HGSOC tissues from the same cohort of patients as the above analysis showed that all five matrisome molecules associated with M0 macrophage phenotype are found predominately in the stroma (Supplemental Figure 5, Supplemental Video 1). Finally, we used an online resource^30^ to determine whether the ECM signature that associates with M0 macrophages correlated with overall survival (OS), where we found a significant decrease in the median OS of ovarian cancer patients (OS = 28.25 months) compared to the low expression cohort (OS = 45.63 months; Supplemental Figure 6A), with a greater hazard ratio (HR = 1.56) than the expression of any gene alone, except COL11A1 or SFRP2 (HR = 1.78 and HR = 1.66, respectively, Supplemental Figure 7). Further analysis across 19 carcinomas found 12 cancers (including ovarian cancer) to have reduced patient OS with high expression of the ECM signature associated with M0 macrophages (Supplemental Figure 6B). Taken together, we found that the composition of the tumor ECM associates most strongly with a M0 macrophage subtype which is poorly characterized in HGSOC, and five tumor ECM proteins which strongly associated with this macrophage subtype showed a reduced cancer cell killing signature and poor prognosis across 12 cancer types.

### Generation of a decellularized tissue model of ovarian cancer metastases

Having found tumor ECM composition associates with a poor prognostic M0 TAM signature (Figure 1), we next evaluated whether this association resulted from the composition of ECM, using a decellularized tissue model of tumor ECM, made from a library of 39 human omental ovarian tumor samples (Supplemental Table 2, Figure 2A). Prior to decellularization, we characterized each tissue for disease content, and immune infiltrate (Supplemental Figure 8). Following characterization, each tissue was cut to a uniform thickness (300 μm) using vibratome sectioning, and decellularized by cell lysis, lipid extraction, and DNA degradation. Tissues were effectively decellularized, shown by nucleic acid content and nuclei content (Figure 2B-C, Supplemental Figure 9). We next assessed the content of four ECM molecules (collagen 1 (COL1A1), FN1, VCAN, and cathepsin-B (CTSB)) and one glycosaminoglycan, chondroitin sulfate (CS), selected based on either their high level of expression in cancer tissues^1^ and/or their association with the macrophage phenotype (Figure 1). We compared the five markers between matched cellularized and decellularized tissues and found expression levels were comparable for four of the five proteins (Figure 2D, Supplemental Figure 10A). CTSB appeared to increase after decellularization which was an artefact resulting from removal of the cellular component, as the presence of hemotoxylin in the cellularized tissue could be overlapping and masking the intensity of CTSB. Chondroitin sulfate (CS), which decorates the surface of many proteoglycans and comprises a major constituent of the post-translational environment, was also used to indicate tissue composition integrity after decellularization (Figure 2D). We next tested whether decellularization affected the ECM architecture by scanning electron microscopy (SEM), by comparing ECM fiber diameter and alignment between matched cellularized and decellularized tissues (Figure 2E, Supplemental Figure 10B) which were comparable for both low and high disease samples (Figure 2F-G). Finally, we tested whether the decellularized tissues were a biocompatible material for cell culture. IF revealed monocytes/ macrophages were viable for a period of 7 days (Figure 2H) and flow cytometry revealed macrophages could be collected after 14 days. However, cell death increased when compared to tissue culture plastic (TCP) (approximately 5-10% more cell death, Figure 2I, Supplemental Figure 10C), resulting from the removal of the cells from the tissue prior to flow cytometry analysis. We next optimized the cell recovery after finding a significant effect on cell marker expression between macrophages isolated from TCP versus decellularized tissue when using enzyme-based dissociation solutions, which could be negated by using an enzyme-free buffer (Supplemental Figure 11). Taken together, we concluded that the decellularization process maintained the ECM architecture with minimal effect on the quantity of major ECM components, and decellularized tissues were a suitable platform for cell culture, where cells could be visualized on the tissue or removed as single cell suspensions for further assays or analysis.

**Figure 2.**
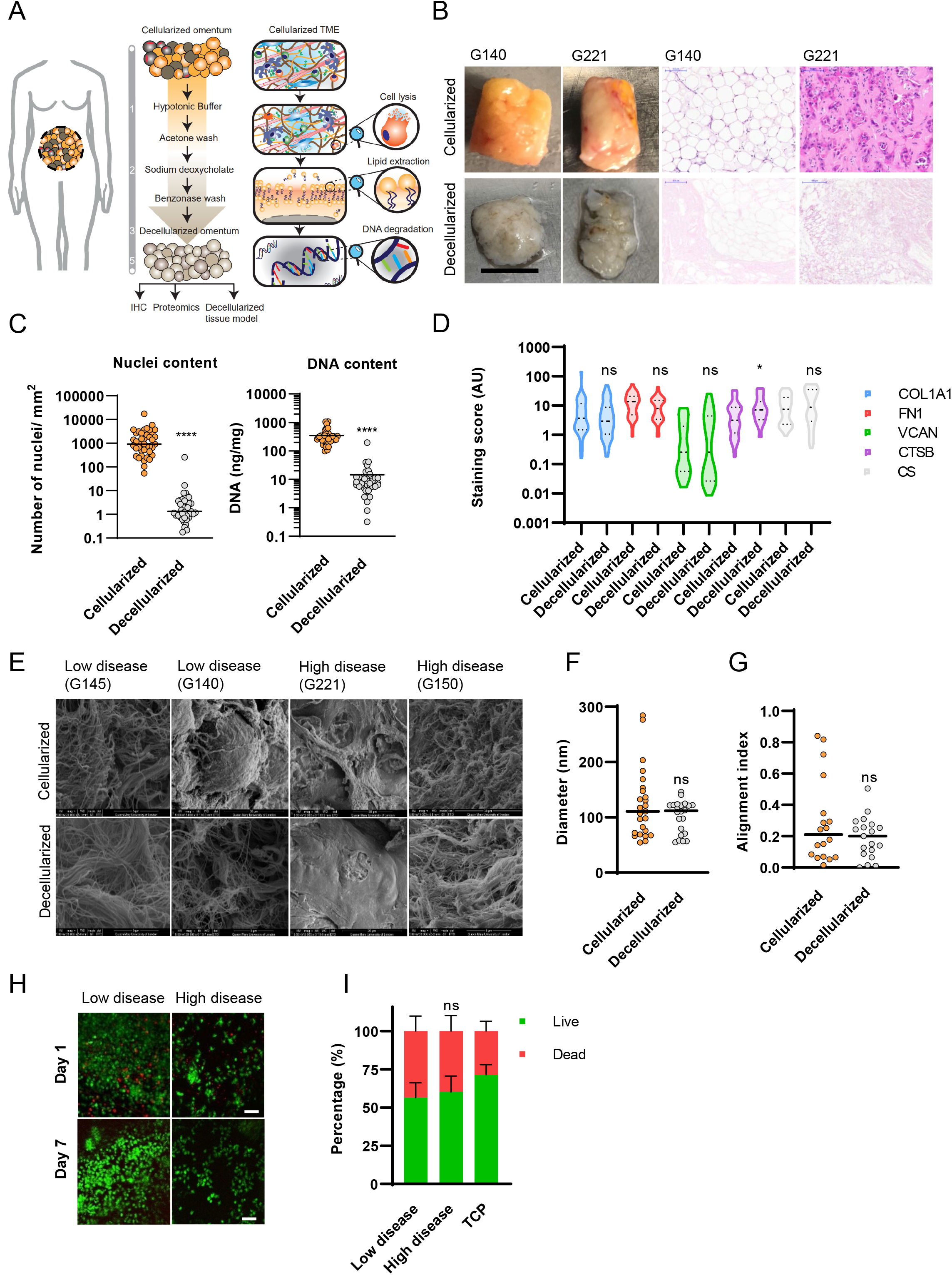
A decellularized tissue model of ovarian cancer metastasis. **A)** Schematic of the decellularization procedure and tissue processing. **B)** Fresh tissue biopsies taken from ovarian cancer patients. Histologically uninvolved samples from benign tumors were used as part of the group of low disease tissues. H&E staining of sectioned ovarian cancer samples. **C)** Nuclei and nucleic acid content in cellularized and decellularized tissue samples (n=39). **D)** IHC staining analysis for matrix molecules in cellularized and decellularized samples. **E)** Representative Scanning Electron Microscopy (SEM) images for cellularized and decellularized samples. **F)** Quantitative fiber diameter analysis from SEM images. Three fields of view were chosen at random in cellularized and matched decellularized tissue micrographs (N = 15). Fiber diameter angles were recorded (minimum 30 fibers quantified per field of view). **G)** Quantitative fiber alignment (alignment index) from SEM images (N = 15). Closer to 1 the more aligned the fibers, closer to 0 the more disorganized the fibers. **H)**Representative live/ dead immunofluorescence (IF) images from decellularized tissue model cultures using monocytes/ macrophages at day 1 and 7. **I)** Bar plot of viable macrophages collected from low disease samples, high disease samples and tissue culture plastic (TCP) (N = 3).

### The M0 associated ECM signature is also a feature within the present tissue library and positively associated with TAM infiltration

We investigated whether the association of ECM composition and TAM phenotype (Figure 1) was a feature in the library of tissues used here to make the decellularized tissue platform (prepared in Figure 2), through integration of proteomics and immune cell counts using IHC data (Figure 3A, immune cell counts per sample are shown in Supplemental Figure 13). We characterized the ECM of decellularized samples using ECM focused proteomics^31^ (Supplemental Figure 14A, and Supplemental Table 3). Proteomics data indicated collagens constituted the largest proportion of the ECM by abundance (35%), followed by glycoproteins (25%) and proteoglycans (25%), ECM-regulators (8%), ECM-affiliated proteins (5%), and finally ECM secreted factors (1%) (Supplemental Figure 14A). We grouped the samples by their ECM composition, which identified five ECM composition groups (ECGs) (Figure 3B). ECGs were compared to the disease score (the sum of the % area of PAX8^+^ cells and % area of stroma) (Supplemental Figure 14B, Supplemental Table 4), and found samples split into two groups corresponding to low disease score (ECG1 and 2), where samples primarily consisted of adipocytes, or high disease score groups (ECG3, 4, and 5), where samples contained high levels of PAX8^+^ cells and stroma (Figure 3B, Supplemental Figure 14B). There was a significant difference in the disease scores between ECG1-2 when compared against ECG3, 4, or 5, indicating the change in ECM composition that occurs as disease progresses^1^. Next, we analyzed the difference in ECM composition between ECG3, 4, and 5. These groups represented tumor tissues with similar disease scores (Supplemental Figure 14B) but different ECM compositions (Figure 3B). Overall expression of major ECM categories between ECGs 3-5 appeared to be similar, with the exception of ECG4, where an expansion of ECM-glycoproteins was observed (Supplemental Figure 14C). Next, we analyzed the differentially expressed (DE) proteins between each ECG and found each had a different pattern of ECM protein expression resulting from changes in the abundance of specific molecules (Figure 3C). We explored if the ECM signature associated with M0 macrophages (Figure 1F) across the ECG groups. Individually, all four molecules tended to be highly expressed in ECG3 and ECG5 and appeared highest in ECG5, with the exception of COL11A1 which did not correlate with a specific ECG group(s). In particular, FN1 expression was significantly higher in ECG5 (Figure 3D). We also compared the ECM signature associated with M0 macrophages (Figure 1F) across the five ECG groups finding ECG5 was enriched for the signature (Figure 3D, right-hand panel). These data indicated ECM composition can vary across ovarian patient samples, an observation we had previously noted from transcriptomic data for primary HGSOC within The Cancer Genome Atlas^1,32^. Taken together, the heterogeneity in ECM composition across tumors with similar tumor and stromal compositions (or disease scores) indicates that the pattern of tumor ECM is independent from the extent of disease. To explore the localization of these matrisome molecules, we integrated the above proteomics data with an analysis which defined adipose, stroma and tumor areas in the same tissues using IHC data. Expression of FN1 and VCAN was significantly positively correlated with the percentage of stroma area, but not tumor area, and were significantly negatively correlated with the percentage of adipose area, indicating that, for the most part, these ECM molecules were being expressed by cells found within the stroma (Supplemental Figure 15). Subsequently, proteomic data of cell-derived matrices (CDMs) from patient-derived omental fibroblasts and metastatic HGSOC malignant cell lines were measured to explore expression of these matrisome molecules in *in vitro* cultures. In line with our previous results, four of the matrisome molecules associated with M0 macrophages (FN1, VCAN, COL11A1, MXRA5) were more highly expressed by patient-derived omental fibroblasts than malignant patient-derived HGSOC cell lines (Supplemental Figure 16). We wondered if the ECM patterns seen in our decellularized tissue library (Figure 3C) may correlate with the immune infiltrate, as we had found from our informatics analysis (Figure 1). We integrated ECM proteomics data (Figure 3C) with immune cell counts (Supplemental Figure 8 and 13, Supplemental Table 5). Immune cell counts were evaluated across ECGs (Figure 3E-F). Low disease omental samples (ECG1-2) composed almost entirely of adipose tissue (Supplemental Figure 8D) and had ECM compositions comparable to normal tissues^31^ had low immune cell abundances (Figure 3E-F). In contrast, the level of immune infiltrates changed significantly between ECGs 3<4<5 (Figure 3E). We found that macrophages (CD68^+^) were the dominant immune cell type in ECG3-5, being highest in ECG5 (Figure 3E). These data confirmed the earlier analysis we had done on a separate set of tissues (Figure 1). The number of CD4^+^ and CD8^+^ cells varied between ECG3-5, with CD8^+^ cells highest in ECG5 and CD4^+^, FOXP3^+^, and CD20^+^ highest in ECG4. We calculated Z-scores using cell count data and categorized by ECG (Figure 3F), which revealed the trend of increasing CD8^+^ and CD68^+^ cell counts between the three groups ECG3<ECG4<ECG5. We evaluated if the presence of tumor cells (PAX8^+^cells) may also correlate with immune cell counts between ECG3-5, however we found no correlation when looking at tumor samples (Supplemental figure 14D-G). Together, these data identified an association between the tumor ECM composition and immune infiltrate that confirmed our previous findings (Figure 1). Next, we investigated whether the ECM composition could educate the M0 TAM phenotype that associated with ECM composition in HGSOC.

**Figure 3.**
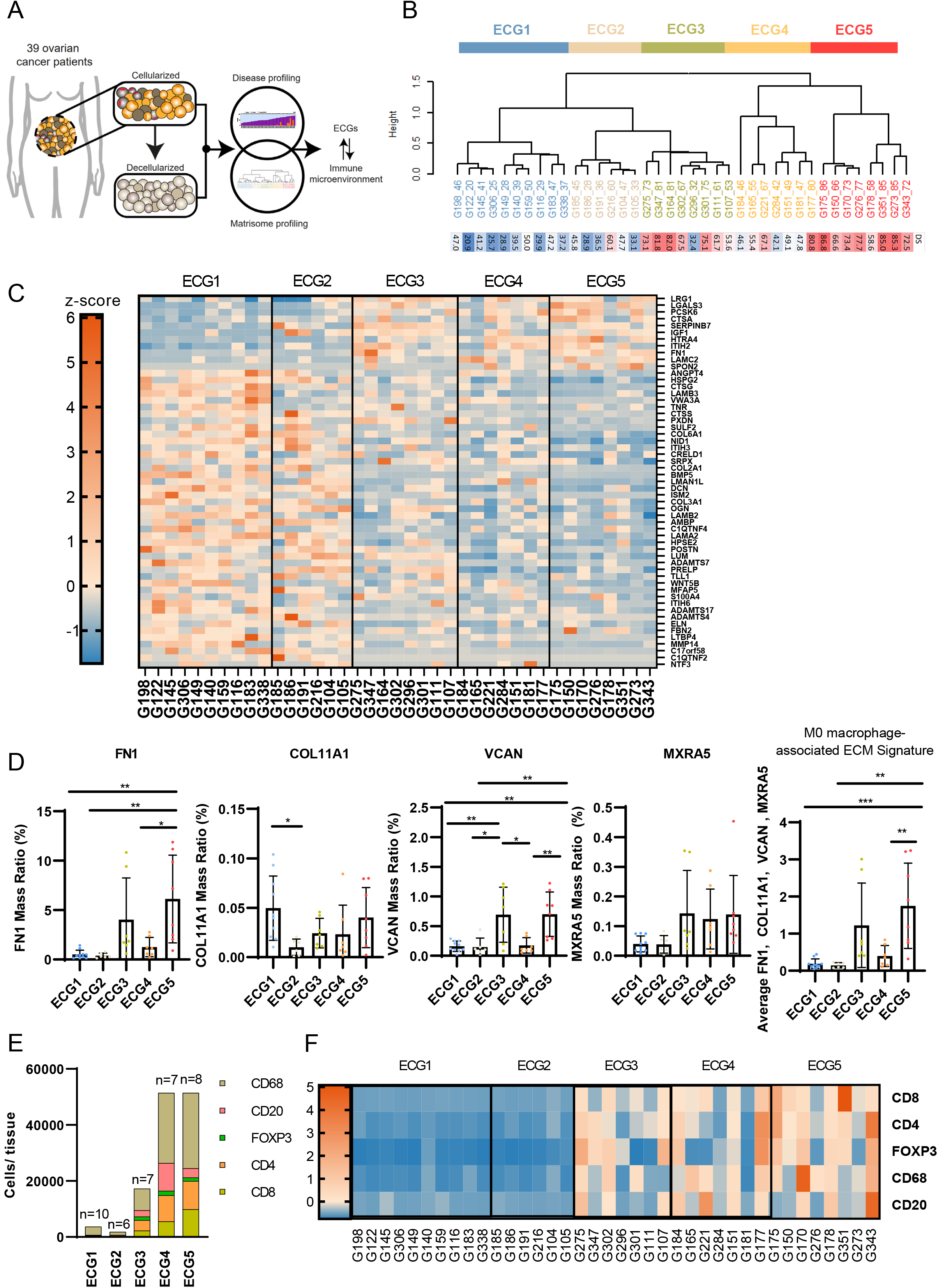
ECM composition correlates with the immune cell landscape. **A)** Flowchart of integration analysis using the disease score (disease profiling) and ECM proteomics (matrisome profiling) of ovarian metastatic samples to define the ECM composition and explore the synergy with ECM composition and the immune cell landscape. **B)** Hierarchal unsupervised clustering (ward.D2 method), separated into ECM composition groups (ECG) 1-5. Labels: tissue ID _disease score. Disease score (DS) displayed as heatmap from low disease score (blue) to high (red). **C)** Heat map of positively and negatively regulated matrisome proteins, columns grouped by ECG1-5. **D)** Bar plots of protein expression for FN1, VCAN, COL11A1 and MXRA5 in ECG1-5. One way ANOVA with Tukey’s post-hoc test, significance between each group is presented as **p < 0.01 and *p < 0.05. **E)** Mean number of cells/ number of tissues between ECG1-5 (N = 36). **F)** Z-scores of IHC immune cells/ mm^2^ (N = 38).

### In vitro ECM educated TAMs share gene profiles with M0 TAMs found in human tissues

Using the decellularized tissue model, we explored whether the tumor ECM could directly educate monocytes to differentiate into the M0 TAM phenotype found in ovarian metastases (Figure 1 and 3). We cultured monocytes derived from healthy donor peripheral blood mononuclear cells (PBMCs) on decellularized tissues from our library (Figure 2 and Supplemental Figure 8) that represented ECM compositions of low ‘adjacent’ to disease tissues (ECG1-2, x4 tissues) or high diseased ‘tumor’ tissues (ECG3 & 5, x4 tissues) (Figure 4A), and after 14 days, macrophages were isolated and bulk RNA-sequencing (RNAseq) was performed (Figure 4B, Supplemental Figure 17A). Transcriptomic profiles grouped samples based on which decellularized tissue they were cultured on (Figure 4B, Supplemental Figure 17B). Differential gene expression analysis identified a total of 3,613 genes between macrophages cultured on the tumor tissue (ECG3 & 5, 1,839 upregulated genes) and macrophages educated on the adjacent tissue (ECG1 & 2, 1,774 upregulated genes) (Supplemental Figure 17C-D). CIBERSORTx analysis of DE genes revealed that macrophages educated by the tumor or adjacent decellularized ECM shared gene profiles with the M0 macrophage subtype (Figure 4C), leading us to term these as extracellular **ma**trix-educated **m**acrophage**s** (MAMs). Gene analysis revealed tumor MAMs contained significantly more of the M0 macrophage subtype and shared signatures with M2-like macrophages, when compared to adjacent MAMs (Figure 4D-E, Supplemental Figure 18). Conversely, adjacent MAMs showed similarities to an M1-like phenotype, when compared to tumor MAMs (Figure 4D-E). Subsequently, we performed an analysis to relate the significantly up-regulated genes in tumor or adjacent MAMs to a spectrum-model^33^ of macrophage activation generated by stimulating human CD14+ monocyte-derived macrophages using 29 discrete stimuli that revealed 49 co-expressed gene modules (Supplemental Figure 19). Tumor MAMs significantly positively correlated with modules that occur upon *in vitro* activation with stimuli linked to fatty acids (palmitic acid, oleic acid, linoleic acid) and to chronic inflammation (TNF, prostaglandin E2, TLR2-ligand; Supplemental Figure 19B). Whereas, adjacent MAMs significantly positively correlate with modules that occur upon *in vitro* activation with stimuli linked to LPS and to chronic inflammation (Supplemental Figure 19B). Neither MAM phenotype is associated with stimuli linked to M1 or M2 polarization, which was in line with the analysis on whole cancer tissues (Figure 1). These data suggest tumor and adjacent MAMs undergo transcriptional reprogramming, however, overall these changes are not consistent with a strict M1-M2-axis re-polarisation as assessed by individual gene expression changes. Taken together, we conclude that tumor MAMs share transcriptomic similarities to the macrophage phenotype seen in HGSOC tissues, and therefore we conclude the ECM is capable of stimulating the TAM population seen in HGSOC.

**Figure 4.**
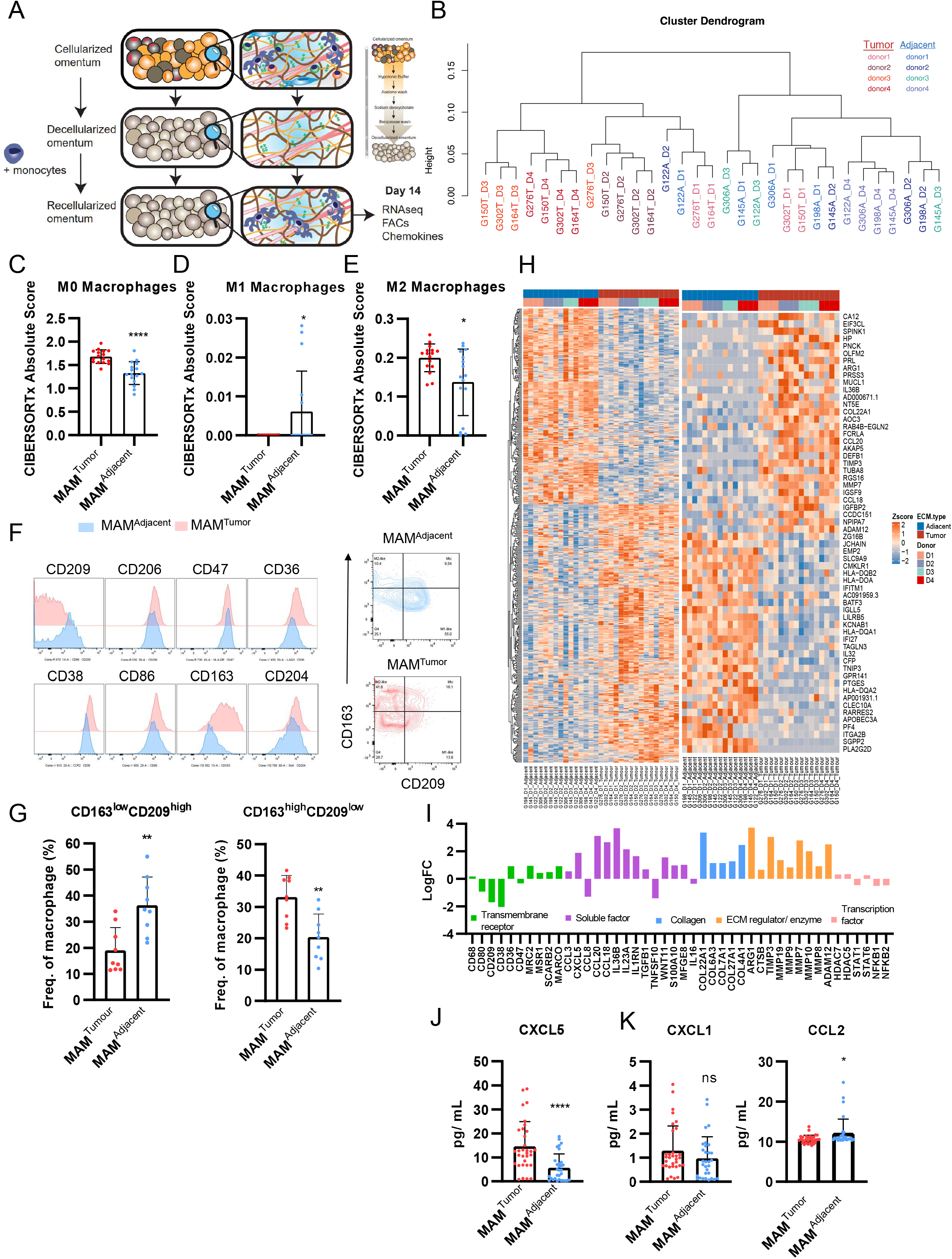
Tumor ECM alters the macrophage transcriptome. **A)** Schematic of macrophage decellularized tissue culture. **B)** Unsupervised cluster dendrogram using DE genes. **C-E)** Bar plots of CIBERSORTx analysis using DE genes between tumor and adjacent MAMs for **C)** M0 macrophage, **D)** M1 macrophage and **E)** M2 macrophage signatures. **F)** Histograms of transmembrane receptors expressed on tumor and adjacent ECM cultured macrophages. Representative contour plot of tumor and adjacent MAMs. **G)** Bar plots of expression pattern of CD163 and CD209 between tumor and adjacent MAMs, from flow cytometry shown as percentage in CD45^+^ CD14^+^ macrophage gating CD163 and CD209. Mann-Whitney U test, p = 0.004 and Unpaired T test, p = 0.0016, respectively (N = 3). **H)** Heatmap of row z-scores of log2TPM gene expression for top 30 up- and downregulated genes. (adj. p < 0.05, logFC > |1|, protein coding). **I)** Bar plot of selected DE genes between tumor versus adjacent cultured macrophages. **J)** LEGENDPLEX™ assay of secreted CXCL5. Mann-Whitney U test, p < 0.0001. **K)** LEGENDPLEX™ assay of secreted CXCL1 and CCL2 expression levels.

### Tumor MAMs have an immunoregulatory and tissue remodeling phenotype

We next investigated the macrophage phenotypic transition by characterizing cell surface markers in MAMs educated by tumor (ECG3&5) or adjacent (ECG1 & 2) decellularized tissue. We used flow cytometry comparing the expression of macrophage markers including CD14, CD11b, CD45, CD163, CD206, CD86, CD38, CD209, CD47, CD36, and CD204, between tumor and adjacent MAMs (Figure 4F, Supplemental Figure 20). Transmembrane receptors including alternatively activated macrophage marker, CD163, were significantly upregulated on tumor MAMs, while mannose receptor, CD209, was upregulated on adjacent MAMs (Figure 4G). These data confirmed our observation from the CIBERSORTx transcriptomic analysis (Figure 4C-E) that the matured macrophages were not predominantly polarized into the classically defined ‘M1’ or ‘M2’-like states. Next, we looked at the total and the top 30 DE genes that associated with tumor MAMs (Figure 4H), which included transmembrane receptors, soluble factors, ECM-collagens, ECM-regulators, transcription factors, scavenger receptors like CD36, C-X-C motif chemokine ligand 5 (CXCL5), arginase (ARG1) and a number of matrix metalloproteinases (MMPs) (Figure 4I). The patterns of upregulated genes indicated tumor MAMs may have an ECM remodeling activity. Since several soluble factors seemed to be expressed by tumor MAMs, we used a multiplex assay (LEGENDplex™) to characterize a panel of macrophage secreted factors at the protein level (Supplemental Figure 21, Supplemental Table 6). This analysis revealed chemokine CXCL5, a chemoattractant of T cells, and a member of the M0 CIBERSORTx signature, was significantly upregulated by tumor MAMs, at gene and protein level (Figure 4I-J). Other secreted factors including CCL2 which was significantly upregulated by adjacent MAMs, and CXCL1, which showed no change between tumor or adjacent MAMs was detected at very low levels (Figure 4K, Supplemental Figure 21). To explore which biological processes may be altered in tumor MAMs, we used weighted gene correlation network analysis (WGCNA), which identified 13 coordinately expressed gene programs associated with distinct biological pathways (Figure 5A). Hierarchal cluster analysis separated the gene programs by their similarity (Figure 5B). Gene programs such as purple, tan and brown were significantly upregulated in tumor MAMs, while pink, red, black and others were downregulated (Figure 5C). Tumor MAMs upregulated programs (purple, tan and brown) were distinctly enriched for integrin receptors (blood coagulation pathway), ECM organization and ECM disassembly and leukocyte cell adhesion, while downregulated gene programs (pink, red, turquoise, yellow and black) included pathways of the inflammatory response such as MHC class II antigen processing and presentation (matching our earlier computational analysis on immunity for tissues with a high ECM signature associated with M0 macrophages, Figure 1G), IFN-γ signaling, and defense response (Figure 5D). This indicated tumor MAMs were associating with an integrin-ECM interaction and were involved in ECM remodeling via both construction (ECM organization) and deconstruction (ECM disassembly). We built a tumor MAM gene signature based on DE genes and TAM-like markers (Figure 5E) and correlated this with ECM receptors, collagen degradation molecules, and classic macrophage activation genes which further associated tumor MAMs with a tissue remodeling phenotype (Figure 5F). Taken together, these data describe the tumor MAM phenotype having some alternatively activated character with immunoregulatory, and tissue remodeling capabilities, which are also predicted characteristics of the M0 macrophage phenotype. This suggests these same macrophage processes may also be driven by tumor ECM *in vivo*. We next focused on whether the tumor MAMs may have immunoregulatory properties as predicted from the transcriptomic profiles.

**Figure 5.**
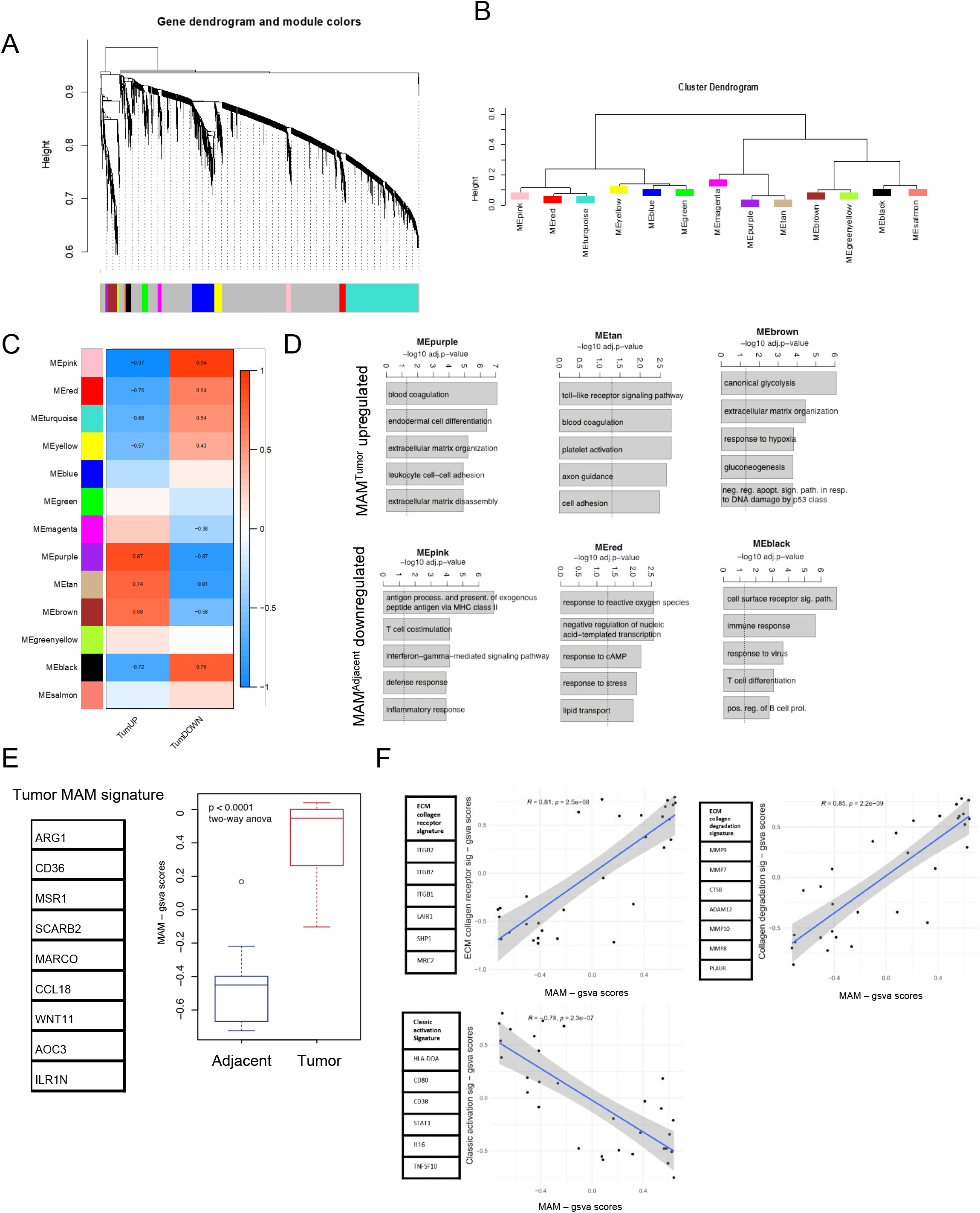
Tumor MAMs infer immunoregulatory phenotype. **A)** WGCNA of human macrophages cultured on tumor and adjacent decellularized tissue showing clusters of co-expressed genes as dendrogram. Colors indicate different modules (gene programs). **B)** Cluster dendrogram of module eigenvalues (MEs) showing associated gene programs. **C)** Heatmap of association of ME with gsva scores of the tumor (vs. adjacent) up and down. Pearson’s r-values are noted for the significant correlations p < 0.05. **D)** Bar plots for significantly enriched gene ontology biological processes in purple, tan, brown, pink, red and black gene programs. Broken line denotes adjusted p = 0.05. **E)** Boxplot of DE Tumor MAM signature genes. **F)** Correlation scatter plots of gsva scores (from gene lists). Correlation coefficient corresponds to Pearson’s correlation.

### Tumor MAMs have reduced phagocytic function and alter T cell activation

Tumor MAMs are predicted to be immunoregulatory based on transcriptomic profiles and their secretome (Figure 4-5). To evaluate their immune function, we first performed a phagocytosis assay, an intrinsic function of macrophages. Tumor MAMs had a reduced phagocytic function compared to adjacent MAMs (Figure 6A). We thought this reduced phagocytosis may be due to the difference in expression of some transmembrane receptors such as CD209 (which was expressed lower in tumor MAMs at gene and protein levels) (Figure 4G, Figure 4I). CD209 has previously correlated with macrophage particle uptake^34^, however we found no difference in the MAMs phagocytic capacity when selecting for CD209^+^ and CD206^+^ macrophages (Figure 6B). Because we previously showed that tumor MAMs have up-regulated toll-like receptor signaling (Figure 5D) which can promote production of pro-inflammatory cytokines and chemokines (Figure 4I), we wondered whether tumor MAMs may affect T cell function. T cells were cultured with low level CD3/ CD28 stimulation with tumor or adjacent MAMs and in parallel, we cultured T cells with tumor or adjacent MAMs conditioned media to test whether these macrophages directly or indirectly (macrophage-secreted factors) modulated T cell phenotype and proliferation (Figure 6C). Tumor MAMs induced inhibitory receptor expression of LAG3 on CD3^+^ T cells, compared to adjacent MAMs and control (Figure 6D). This increase in LAG3 expression was not induced in tumor MAM conditioned media (Figure 6E), indicating a cell-cell interaction between tumor MAM and T cells was needed to induce receptor expression. Upregulation of PD1 and TIM3 on CD3^+^ T cells was also induced by tumor MAMs compared to control and adjacent MAMs, although the difference was not statistically significant in the latter. T cell proliferation was significantly increased by both tumor or adjacent MAMs compared to control (Figure 6F). In contrast, when using MAM conditioned media no difference in T cell proliferation, or activation markers were recorded (Figure 6G). Taken together, *in vitro* co-culture assays support the transcriptomic analysis of tumor MAMs indicating a role for these tumor MAMs in regulating infiltrating T cells within the TME.

**Figure 6.**
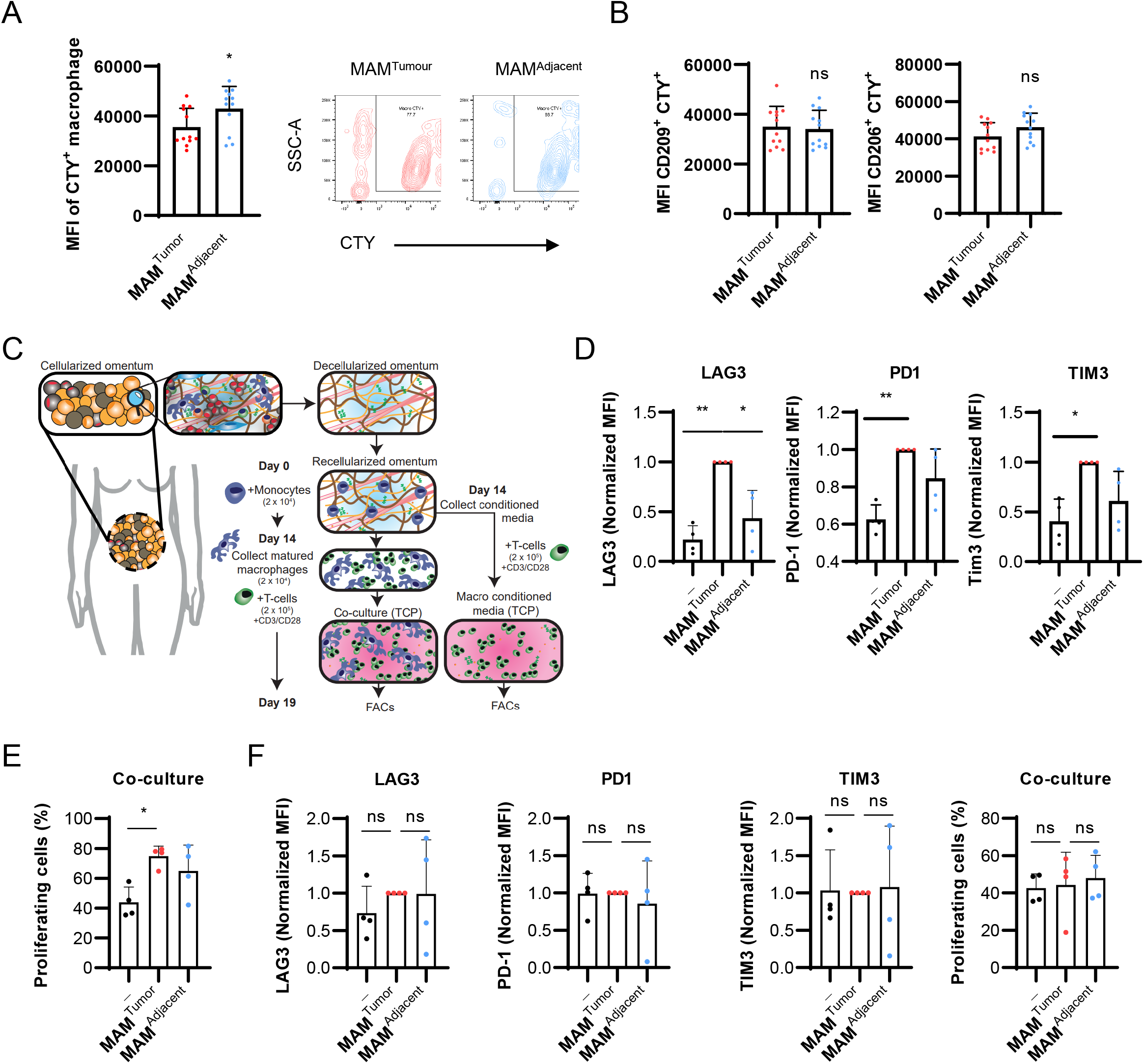
Tumor MAMs have an increased phagocytic response and alter T cell activation in the presence of CD3 and CD28 stimulation. **A)** Flow cytometry analysis of mean fluorescence intensity (MFI) of tumor and adjacent MAMs. p=0.02. Mann-Whitney U test used. Representative contour plot of CTY macrophages cultured between tumor and adjacent ECM. **B)** MFI of CD209 and CD206 CTY cells. P= 0.77, P=0.11, respectively. Unpaired T-test used. n=4. **C)** Schematic of MAMs and T cell co-culture workflow. **D-E)** Normalized MFI of LAG3, PD1 and TIM3 expression on T cells after 5 days co-culture with tumor or adjacent MAM **(D)** or macrophage conditioned media **(E)**. **P<0.01, *P<0.05, one-way ANOVA followed by Dunnett’s multiple comparisons test. **F)** Percentage of proliferating T cells as assessed by CTV dilution after 5 days co-culture with tumor or adjacent MAM (top panel) or macrophage conditioned media (bottom panel). *P<0.05, one-way ANOVA followed by Dunnett’s multiple comparisons test.

## Discussion

The expanding repertoire of biological cancer therapies in recent years has highlighted the influence of the tumor microenvironment in host and therapy response particularly in carcinomas, and significant evidence has linked the composition of the tumor ECM with anti-tumor immunity and immunotherapy responses. There is a growing awareness that the ECM may have a direct role in tumor progression and host immunity, and therefore we set out to investigate whether the ECM directly educates the immune infiltrate. In this study, we show the tumor ECM composition in ovarian metastases and 15 other solid cancers associate with an M0 macrophage phenotype. This ECM composition is largely contributed from fibroblast, however tumor cells are also producing these molecules, in some cases to similar levels seen in fibroblasts. Using a decellularized tissue model of ovarian omental metastases, where the composition and structure of the tumor ECM is maintained, we show that the ECM can directly educate macrophages that overlap with the transcriptomic profile of a prominent TAM population found in HGSOC. We suggest that the reason these TAMs associate with tumor ECM is because they are generated through an interaction of infiltrating monocytes with the tumor ECM, that we termed extracellular matrix-educated macrophages (MAMs). Of note, we found that decellularized tissue alone is sufficient to mature monocytes into macrophages, an observation that is supported by a study of monocytes cultured on a decellularized tissue model of colorectal cancer^35^. Macrophages educated on decellularized tissue undergo a phenotypic transition, shown by different patterns of marker expression and transcriptomic profiles, that is dependent on ECM composition. Macrophages educated by tumor ECM have a TAM phenotype with significantly altered cell surface expression similar to alternatively activated macrophages and a transcriptomic signature comprised of immunoregulatory and tissue remodeling effectors including ARG1, CD36, MSR1, and MARCO. Tumor MAMs have an immunoregulatory effect on T cells in their ability to alter the expression of activation/exhaustion markers on T cells. In co-culture studies, we found tumor MAMs upregulated classical markers of activation on T cells via direct cell-cell interaction. In the present study the mechanism of how ECM educates the macrophage phenotype seen has not been investigated, however transcriptomic data points to two potential mechanisms, both receptor-ECM interactions: There are leukocyte associated Ig-like receptor 1 (LAIR1) which may signal immunosuppressive gene programs when activated through collagen binidng^36^, or sialoglycan-siglec signaling^37^ where sialoglycans upregulated within tumor tissue may be ligands for immune-inhibitory siglecs on infiltrating macrophages. ECM education of some populations of TAMs is supported by other work which has shown tumor-associated glycoproteins like tenascin-C^38^ and the mucin glycoform MUC1-ST^37^ can induce TAM-like phenotypes and inhibit the function of other immune cells. Tissue stiffness has also been implicated in driving macrophage phenotype and based on our previous studies ovarian metastatic tumor tissue is stiffer (10-20kPa) than low disease tissue (0.1-1kPa). Our preliminary investigations indicate decellularization does not have a significant impact on tissue stiffness, and within a synthetic model the substrate stiffness alone is not driving the macrophage phenotype discussed here. The data presented here are supportive of the hypothesis that tumor ECM has a direct effect on tumor immunity. In the present study we have concentrated on macrophages, however there are other immune cell - ECM composition associations within the analysis presented in Figure 1. Whether the ECM is also capable of driving certain phenotypes in these immune cells has not been tested, however we suggest the decellularized tissue model would be suitable to test hypotheses involving other immune cells. The decellularized model allows the cellular component of the TME, which has been documented to stimulate macrophages, to be removed from the ECM in order to study direct ECM effects on macrophages, however it may also be suitable for multi-cell type studies as well as allowing the cellular aspects of the TME to be recapitulated in as little or much detail as required. As more attention continues to build on therapeutic targeting of the tumor microenvironment, we suggest the decellularized tissue model presented here is a useful research tool to help understand the challenges presented by the TME and may help support the development of the next cancer therapies.

## Methods

### Computational integration of immune signatures with tissue composition

First, we conducted a Spearman correlative analysis between disease score defined as percentage of tissue area occupied by malignant cells and stroma, a previously reported tumor ECM signature termed as ‘matrix index’ and the CIBERSORTx or xCell immune signatures from the RNA-seq data from 32 HGSOC patient tissues^1^. This analysis identified M0 macrophages as significantly positively correlated with extent of disease and tumor matrisome. We next looked to see if we could elucidate specific interactions between immune populations and individual ECM molecules. To do this, we utilized a proteomic pipeline categorizing all core ECM proteins and associated molecules to identify 145 matrisome molecules found in both transcriptomic and proteomic datasets^1^. We then performed a Spearman correlative analysis to integrate the CIBERSORTx and xCell immune signatures with patient matched matrisome transcriptomic and proteomic datasets to identify significant immune-gene and immune-protein associations. Five ECM molecules were found in this analysis to be significantly positively associated with disease score and the M0 macrophage signature using both CIBERSORTx and xCell approaches at both the transcriptomic and proteomic level (FN1, VCAN, MXRA5, COL11A1, SFRP2).

### Tissue samples

Patient-derived tissue specimens were resected from ovarian cancer patients at London based hospitals and stored within the REC & HTA-approved Barts Gynae Tissue Bank, HTA license number 12199. REC no: 10/H0304/14 with full written content. Patient information is detailed in Supplemental Table 2.

### Tissue decellularization

Decellularization involved a series of washes first using hypotonic buffer (10mM Tris, 5mM EDTA, 0.1mM phenylmethylsulfonyl fluoride (PMSF) (Sigma-Aldrich, P7626-250MG), pH 8) for 4 hours at room temperature (RT), 100% acetone (Sigma-Aldrich, 179124) 16-18 hours at 4°C, 2.5mM or 4% Sodium deoxycholate (Sigma-Aldrich, D6750-100G) for 4 hours at 4°C, for adjacent or diseased, respectively and nucleic degradation solution (50mM Tris, 1mM Magnesium chloride (MgCl_2_) (Sigma-Aldrich, M8266-100G), 0.1% BSA and 40 units/mL Benzonase nuclease (Sigma-Aldrich, E1014-5KU), pH 8) for 20 hours at 37°C. PBS 1% P/S washes were used between each buffer and at 4°C for 2 days prior to tissue use for *in vitro* cultures. DNeasy^®^ Blood and Tissue kit (Qiagen, 69504) following manufacturer’s instructions. Nucleic acid (DNA content) was quantified using a nanodrop 2000 spectrophotometer (Thermofisher). H&E stains section were quantified using QuPath tissue imaging software that used a cell detection system to determine total cell number. DNA and nuclei content data provided in Supplemental Table 3.

### Tissue proteomics

The ECM component was enriched from frozen whole decellularized tissue sections (20 x 30 μm sections, approximately 40-50 mg of tissue) and analyzed as previously described^31^. Briefly, the extracted proteins were reduced, alkylated and digested with trypsin. Peptides were separated by nanoflow ultra-high pressure liquid chromatography (UPLC, NanoAcquity, Waters) and analyzed by mass spectrometry using a LTQ-Orbitrap XL mass spectrometer (Thermo Fisher Scientific). Proteomics data provided in Supplemental Table 4.

### Decellularized tissue macrophage culture

Sliced decellularized tissues were equilibrated with DMEM 1% P/S (no serum) at 4°C overnight. Decellularized tissues were removed from DMEM 1% P/S and excess liquid removed. Decellularized tissue slices were placed into the wells of 96-well plate. To the decellularized tissues, 25μL of isolated monocytes were added to the centre of the tissue at seeding density of 2 x 10^5^ / 25μL. The monocytes were incubated with the decellularized tissue at 37°C for 2 hours to allow the cells to attach to the tissue. After 2 hours, 200μL DMEM supplemented with 10% human AB serum (HS) (Sigma-Aldrich) and 1% P/S was added carefully to the cultures, not to disturb the tissue adhered cells. Cultures were maintained for 14 days.

### Phagocytosis assay

K562 cells were collected from a T75 flask and then stained with CellTrace yellow (CTY) in PBS, 1:10,000 for 20 minutes at 37°C then 20mL RPMI was added and incubated 5 minutes at 37°C. CTY^+^ K562 cells were centrifuged and resuspended in RPMI at 5 × 10^5^ cells/mL and plated onto an ultra-low attachment plate at 25,000 cells/well and incubated with anti-CD47 antibody at 40μg/mL for 1 hour at 37°C. In the meantime, TEMs were collected from decellularized tissues and seeded into a fresh ultra-low attachment plate at 10,000 cells/ well. K562 cells/ well were washed in RPMI, resuspended in 100μL RPMI and combined with TEMs for 2 hours at 37°C. Cells were then washed and stained with a flow cytometry antibody cocktail and analysed by flow cytometry. Staining panel used: CD206 (FITC, 1:100), CD209 (APC, 1:100), HLA-DR (AF700, 1:100), CD36 (BV421, 1:100), CD38 (BV605, 1:150), CD86 (BV650, 1:150), CD11b (AF594, 1:200), CD14 (PE-Cy5, 1:150), CD204 (PE-Cy7, 1:100) and Zombie NIR (1:1000) (Biolegend); CTY was captured on the PE channel.

### T cell assay

Human T cells were isolated from frozen PBMCs of autologous samples via the EasySep Human T cell isolation kit (StemCell) according to the manufacturer’s instructions. T cells were then resuspended at 2 × 10^6^ cells/ml in PBS and stained with Cell Trace Violet (1:2000, Invitrogen) for 20 minutes at 37°C. Cells were then washed twice in complete medium (RPMI supplemented with 10%FBS and 1% Penicillin-Streptomycin), seeded in a 96-well plate at 2 × 10^5^ cells per well either alone or in presence of 20 × 10^4^ tumor or adjacent MAMs and were activated with 1 μg/ml anti-CD3 and 5 μg/ml anti-CD28. At day five FACs analysis was performed to evaluate T cell activation and proliferation.

### Laser capture microscopy

Frozen tissue sections from human omental metastasis (14 micron) were cut on to LCM membrane RNAse free slides (previously activated under UV light for 30min). Tissue sections were stained with hematoxylin (a few drops of hematoxylin was added to each section which was then immediately washed in DI water (a few dips), then tap water (a few dips), then submerged in 70%EtOH (30sec), then 100% EtOH (1 min), and finally xylene (30sec) before air drying). Slides were kept on dry-ice where possible, and prior to laser capture. Tumor and stromal areas were then captured using a PALM laser capture microscope (examples of areas taken are shown in Figure S5). Total cut time per slide was 30min max. Cut pieces were pooled from 3-6 tissue sections, and total RNA isolated using Qiagen MinElute columns including on column DNAse treatment as per the manufacturers protocol. Samples sent for sequencing had RNA concentrations in the region of 1800 – 18000 pg/μL and RIN numbers >7.

### Cell-derived matrices (CDMs)

In order to decellularize *in vitro* cultured cell lines, media from 14-day confluent cells (cultured with 50 μg/ml of L-ascorbic acid 2-phosphate (AA2P), dissolved in dH2O)) was first removed and cells were washed using PBS. Cells were then incubated with 37°C warmed extraction buffer (20 mM NH4OH, 0.5% Triton X-100 in PBS) for 20 minutes or until cells showed signs of lysis (visualised by light microscope). Cells were next washed with PBS 70 several times to remove cell bodies. To degrade the cellular DNA, 5-8mL of DNase I (SigmaAldrich) at 10 μg/mL in PBS was incubated with cells at 37°C for 2 hours. Following this, cells were washed with PBS 1% P/S several times. The resulting CDMs were stored at −80°C.

