## Supplemental Methods for "Extracellular matrix educates a tumor macrophage phenotype found in ovarian cancer metastasis"

*Study approval.* Ovarian cancer patient samples were kindly donated by women undergoing surgery at Barts Health NHS trust between 2010 and 2017. All tissue obtained was deemed by a pathologist to be surplus to diagnostic and therapeutic requirements and was collected under the terms of Barts Tissue Bank (HTA license number 12199. REC no: 10/H0304/14). Each patient gave written informed consent. The work was conducted in accordance with the Declaration of Helsinki and International Ethical Guidelines for Biomedical Research Involving Human Subjects (CIOMS).

*Tissue samples.* Tissues from ovarian cancer metastasis were resected at London based hospitals and stored within the REC & HTA-approved Barts Gynae Tissue Bank, HTA licence number 12199. REC no: 10/H0304/14 with full written content.

*Blood samples.* Use of anonymised blood samples obtained from Cambridge Biosciences was covered by ethical review board (IRB) approval and informed consent. Approved uses include the isolation of white blood cells for scientific and medical research, including genetic analysis, at a university or pharmaceutical company.

*Data collection.* The raw HGSOC count per million (CPM) RNA-seq dataset (GSE71340; (9)) collected in a previous study^1^, was downloaded from the NCBI Gene Expression Omnibus (GEO) database^2^ and protein mass ratios (PXD004060^1^) from the PRIDE database^3^. Corresponding HGSOC cellularity data, obtained by manually counting six major leukocyte subtypes was obtained from the CanBuild site (canbuild.org.uk^1^). While 36 patient samples were initially obtained, only 32 samples have all 3 corresponding datasets.

*Deconvolution of bulk RNA-seq profiles.* Log2 base RNA-seq data was delogged and processed to fit the input requirements of each computational method. CIBERSORT and CIBERSORTx estimations were performed against the standard LM22 signature gene file in absolute mode with 100 permutations and quantile normalisation disabled. For CIBERSORTx, impute cell fractions analysis module was selected in custom mode, with B-mode batch correction applied using LM22 source GEP. xCell analysis was performed using the N = 64 gene signature and the RNA-seq box ticked.

*Merging cell subtype estimates for correlation to IHC marker cell counts***.** From the default cell type estimations by CIBERSORT and xCell, we identified all cell types that express each of the 6 immune markers for which cell count data was available (CD3, CD4, CD8, CD45RO, FOXP3, CD68). CD3 cell scores are calculated as the sum of CD8 T cells, CD4 naïve T cells, 𝛾𝛿 T cells, CD4 memory T cells and follicular helper T cells (CIBERSORT only) abundances; CD4 cell scores the sum of CD4 naïve T cells, T regs, CD4 memory cells, follicular helper T cells (CIBERSORT only), 𝛾𝛿 T cells, in addition to CD4 Tem and CD4 Tcm for xCell only; CD8 cell scores as just CD8 T cells for CIBERSORT, in addition to CD8 Tcm, CD8 Tem and CD8 naïve T cells for xCell. CD68 cell scores are calculated as the sum of monocytes, macrophages M0, M1 and M2, and dendritic cells (no immature DC); CD45RO just CD4 memory T cells and FOXP3 just Tregs. Spearman’s correlation analysis between the marker cell count data and the marker cell estimation scores was then performed.

*Clinical outcome analysis.* Delogged HGSOC RNA-seq data was input to estimate the tumor progression within the cancer immunity cycle, using the tracking tumor immunophenotype (TIP) meta-server^4^. Samples were separated into the top 12 high and bottom 12 low expressing samples. Kaplan-Meier plots, hazard ratios and log ranked p significance values were generated using gene expression omnibus (GEO), European genome archive (EGA) and the cancer genome atlas (TCGA) RNA-seq databases, via the KM plotter online meta-analysis tool^5^.

*Statistic and bioinformatics analysis***.** All statistical analysis was performed and graphics drawn using either GraphPad Prism (Version 8.4.3) or the statistical programming language RStudio (version3) using the following software plugins: Hmisc for correlation analysis, gplots for correlation scatter plots, ggplot2 for bar charts, pheatmap for heatmaps, dendextend for dendrograms and ggpubr for editing figures to publication standard. Multivariate correlations were calculated using Spearman’s or Pearson’s correlation as appropriate, applied on linear or log transformed data, where p < 0.05 is considered significant unless otherwise specified and indicated with asterisk: *p < 0.05, **p < 0.01, ***p < 0.005.

*RNA isolation and sequencing.* Total RNA was extracted from macrophage cultures on decellularized tissue using RLT buffer (Qiagen) and rigorously vortexed. Samples were processed using the RNeasy Micro Kit (Qiagen) following manufacturer’s instructions. RNA quality and integrity assessments were performed at Oxford Genomics. RNA sequencing was performed at Oxford Genomics. Material was quantified using RiboGreen (Invitrogen) on the FLUOstar OPTIMA plate reader (BMG Labtech) and the size profile and integrity analysed on the 2200 or 4200 TapeStation (Agilent, RNA ScreenTape). RIN estimates (where available) were between 7 and 9.7. Input material was normalised to 10 ng (or maximum mass available) prior to library preparation. Polyadenylated transcript enrichment and strand specific library preparation was completed using NEBNext Ultra II mRNA kit (NEB) following manufacturer’s instructions. Libraries were amplified (18 cycles) on a Tetrad (Bio-Rad) using in-house unique dual indexing primers (based on DOI: 10.1186/1472-6750-13-104). Individual libraries were normalised using Qubit, and the size profile was analysed on the 2200 or 4200 TapeStation. Individual libraries were normalised and pooled together accordingly. The pooled library was diluted to ~10 nM for storage. The 10 nM library was denatured and further diluted prior to loading on the sequencer. Paired end sequencing was performed using a NovaSeq6000 platform (Illumina, NovaSeq 6000 S2/S4 reagent kit v1.5, 300 cycles), generating a raw read count of >23 million reads per sample.

*Cytokine and chemokine analysis.* Cytokine and chemokines were assayed using LEGENDPLEX™ platform according to manufacturer’s instructions. The human proinflammatory chemokine panel 1 kit (cat: 741081) was used to quantitatively measure the levels of 12 human chemokines from macrophage conditioned media. Chemokines included CXCL8, CXCL10, CCL11, CCL17, MCP-1, CCL3, CXCL9, CXCL5, CCL20, CXCL1, CXCL11, CCL4. A total of 32 conditions were analysed in duplicates and followed manufacturer’s instructions. The Biolegend cloud-based data analysis software suite was used to analyse flow cytometry data. Top standards were set based on manufacturers guide for that specific lot number kit (B326082) and were: CXCL8 (21ng/mL), CXCL10 (11ng/mL), CCL11 (12ng/mL), CCL17 (10ng/mL), MCP-1 (13ng/mL), CCL3 (32ng/mL), CXCL9 (4ng/mL), CXCL5 (6ng/mL), CCL20 (2ng/mL), CXCL1 (4ng/mL), CXCL11 (4ng/mL), CCL4 (3ng/mL).

*Histochemical analysis.* Frozen tissues were fixed in in 4 % paraformaldehyde (PFA) and cryosectioned to 8-10 µm slices. All tissue sections were scanned using a 3DHISTECH Panoramic 250 digital slide scanner (3DHISTECH, Hungary) and the resulting scans were analysed using Definiens software (Definiens AG, Germany). Disease scores were determined firstly by manually defining regions of interest in the tissue that represented tumor (PAX8), stroma, and fat (adipocytes) and then training the software to recognize these regions of interest. Disease score was expressed as a percentage of the whole tissue area that contained tumor and/or stroma (Figure 2B-D). Paraffin embedded tissues were submerged in xylene and then a series of ethanol washes of decreasing concentration for 2 x 2 min each (100 %, 90 %, 70 %, and 50 %). Antigen retrieval was performed for 10 min using vector antigen unmasking buffer and a pressure cooker. Tissue sections were then washed with DAKO wash buffer followed by application of H2O2 for 5 min. Blocking was performed using 5 % BSA for 20 min at RT followed by incubation with primary antibody in biogenex antibody diluent for 30 min. After 3 x washes, biogenex super enhancer was added for 20 min and then washed off before addition of biogenex ss label poly-HRP for 30 min. Tissues were washed three times before addition of DAB chromagen for 3 min followed by washing to stop further DAB development. Tissues were counterstained with haematoxylin followed by washing with H2O and ethanol solutions of increasing concentration for 2 min each (50 %, 70 %, 90 %, 100 %) and then 2 x xylene. Samples were then mounted and scanned using the 3DHISTECH Panoramic digital slide scanner. Immune cells were counted using QuPath.

*Matrix staining*. Immunohistochemical staining for ECM proteins was performed on 4 µm slides of FFPE human omentum tissue as described above.

*Antibodies*. The following antibodies were used for immunohistochemical analyses: anti-PAX8 (clone BC12, ab124445; 1:1000), anti-FOXP3 (clone eBio7979 (221D/D3), 14-7979-82; 1:1500) from Invitrogen, UK; anti-CD4 (clone 4B12, M7310; 1:300), anti-CD8 (clone C8/144B, ab75129; 1:500), anti-CD68 (clone PG-M1; 1:12000), anti-CD163 (clone 10D6, MA5-11458; 1:1000), anti-CD20cy (clone ), anti-VCAN (polyclonal, Ab202906; 1:200), anti-COL1A1 (polyclonal, HPA011795; 1:500), anti-FN1 (polyclonal, Ab23750; 1:500), anti-CTSB (ab125067; 1:50), from Abcam, anti-CS (clone CS-56, Ab11570; 1:600).

*Flow cytometry analysis.* Macrophages were initially washed using PBS and then stained with 1:1000 Zombie™ NIR in PBS supplemented with Fc receptor and monocyte blocker (Biolegend) for 20 minutes at RT. Cells were then washed and stained in 50μL total volume PBS, 2% BSA, 2mM EDTA (FACS buffer) using an antibody master mix for 20 minutes at 4°C. Subsequent stained cells were washed in FACS buffer and analysed. Monocyte derived macrophage staining panel used: CD206 (FITC, 1:100), CD45 (PerCP, 1:100), CD209 (APC, 1:100), CD47 (AF700, 1:100), CD36 (BV421, 1:100), CD38 (BV605, 1:100), CD86 (BV650, 1:100), CD163 (PE, 1:100), CD11b (AF594, 1:100), CD14 (PE-Cy5, 1:100), and CD204 (PE-Cy7, 1:100) (Biolegend). Cytofluorimetric analysis was performed using a 4-laser Fortessa flow cytometer (BD). For macrophages 2,000 live events were collected and selected by Zombie NIR gating strategy. Unstained, Zombie™ stained cells spiked with unstained cells and fluorescence minus one (FMO) for each antibody in the panel were used as controls to set up cell gating strategies. Compensation control beads (UltraComp eBeads™ compensation beads, Thermofisher; 01-2222-42) were made for each antibody marker and were used to calculate the overlap of fluorescence between laser channels and automatically calculated and applied to each full stain sample. This panel of markers was validated using a titration of antibodies at various concentrations and staining monocytes at day 0 and differentiated macrophages at day 7 and 14. Data was analysed using Flowjo. *T- Cell flow cytometry.* T-cells were harvested and preincubated with the anti-human Fc-Receptor binding inhibitor (Invitrogen) following the manufacture’s instruction. Cells were then stained for 30 minutes on ice using monoclonal antibodies to the following human proteins: CD223-FITC (3DS223H, ThermoFisher), CD8-APC (SK1, Biolegend), CD366-APCCy7 (F38-2E2, Biolegend), CD4-BV711 (SK3, BD Bioscience), CD3-PE (UCHT1, Biolegend) and CD279-PECy7 (EH12.1, BD Bioscience). Viability was tested using the Zombie Aqua Fixable dye (Biolegend). Samples were acquired on LSRFortessa I (Becton Dickinson) and analysed using FlowJo v10.7 (BD FlowJo LLC).

*Disease score quantification.* The level of disease was calculated via IHC analysis of whole tissue sections as a disease score (Supplemental Figure 8B-D). The disease score was devised by a scoring system that quantified the sum of the area of tumor cells (in this case we used PAX8 as marker of ovarian cancer cells) and tissue stroma (Supplemental Figure 8B-C). The tissue stroma was the area of desmoplasia (or tumor ECM) resulting from the invasive tumor within the tissue. Whole tissue sections were analyzed using Definiens® digital image software, which calculated the percentage of the tumor, stroma and adipose content within the tissues (Supplemental Figure 8C-D). The 39 samples were clinically graded as stage III-IV and were collected predominantly from patients with HGSOC, but also patients with high grade clear cell carcinoma, malignant Brenner tumor, Borderline serous cystadenoma, benign fibroma, benign mucinous cystadenoma, sero-mucinous borderline tumor and mucinous adenocarcinoma (Supplemental Table 2 and Supplemental Figure 12). QuPath was used to quantify PAX8+ cells (Supplemental Figure 8E) and PAX8+ cell counts positively correlated with the disease score (Supplemental Figure 8F), indicating the quantity of tumor cells associated with the level of ECM remodeling. The clinical staging of samples did not correlate with the tissue disease score. We used the disease score to account for differences between tissues that may result from the level of disease present.

*Statistical analysis.* All statistical analyses and graphics were performed in the statistical programming language R (version 3.1.3).
