## Supplemental Figure Legends for "Extracellular matrix educates a tumor macrophage phenotype found in ovarian cancer metastasis"

**Supplemental Figure 1. Validation of computational algorithms predicting transcriptomic immune landscape in HGSOC. A-C)** Stacked bar charts of immune cell predictions by CIBERSORT, CIBERSORTx, and xCell respectively, ordered by sample disease score. Cell color codes are presented in a key below with color matching across methods. **D-E)** Scatter plots of **D)** CIBERSORTx and CIBERSORT or **E)** CIBERSORTx and xCell correlations with previous immunohistochemistry staining cell counts using six immune markers present in CANBUILD dataset, in human HGSOC tissues. Regression analysis of summed abundances of all possible predicted immune cell types with each marker was completed to generate the plotted r values. Points are colored based on the significance of their r values from each technique, where p < 0.05 is significant. (Spearman’s correlation, N = 32).

**Supplemental Figure 2. M0 macrophages significantly correlate with MI and disease score. A)** Scatter plot of immune cell estimates using CIBERSORTx, correlated to sample disease scores (DS) and matrix index (MI). Spearman’s regression analysis was completed to generate the plotted r values. Points are colored based on the significance of their r values as depicted in the key, where p<0.05 is significant. N=32 HGSOC samples. **B)** Scatter plots of immune cell estimates using xCell, correlated to sample disease scores (DS) and matrix index (MI). Spearman’s regression analysis was completed to generate the plotted r values. Points are colored based on the significance of their r values as depicted in the key, where p<0.05 is significant. N = 32 HGSOC samples.

**Supplemental Figure 3. M0 macrophages significantly correlate with a matrisome signature at the transcriptomic level for both CIBERSORTx and xCell. A-B)** Heatmap of ECM genes significantly associated with significant immune cells from **A)** CIBERSORTx and **B)** xCell analysis. Spearman’s r values representing correlation between immune abundances and gene expression levels are plotted according to the color scale, with positive correlations in red and negative in blue. Cream colored associations are insignificant with a correlation p > 0.05, N = 32 HGSOC samples. Ordered by unsupervised clustering.

**Supplemental Figure 4.** **Identification of significant ECM proteins associated to disease score.** Barplot illustrates Pearson *p*-values, FDR corrected using the Benjamini & Hochberg method. The dotted line specifies the significance cut-off p = 0.2. All proteins listed were previously identified to have significant associations with immune subsets, at both the gene and protein level. * indicates proteins forming part of the HGSOC matrix index.

**Supplemental Figure 5. Laser capture demonstrates enrichment for matrisome molecules associated with M0 macrophages in the stroma.** Representative images of tissue with areas marked for **A)** tumor and **B)** stroma laser capture**. C)** Validation of tumor and stroma area laser capture using known malignant cell and fibroblast markers, e.g. malignant cell markers: PAX8, EPCAM, and fibroblast marker: ACTA2. **D)** Tumor/Stroma ratio for matrisome proteins associated with M0 macrophages. N=2 (G33, G75).

**Supplemental Figure 6. M0 macrophage ECM signature associates with poorer prognosis in HGSOC and across multiple cancer types. A)** Kaplan-Meier survival curves with overall survival, divided by low and high gene expression levels of the averaged ECM signature associated with M0 macrophages. Serous ovarian cancer patients N = 523. Median survival time is calculated at 50% survival probability. **B)** Multivariate hazard ratio (HR) with 95% CI, derived using multiple datasets across a range of cancer types, with patients divided by low and high gene expression levels of the M0 macrophage-associated ECM signature averaged. HR > 1 indicates that the M0 macrophage-associated ECM signature gene expression is inversely correlated with OS, while HR < 1 shows positively correlated OS. Log ranked p-values significances are presented by asterisks; **** p < 0.0001, ***p < 0.001, **p < 0.01 and *p < 0.05. KIRP N = 288, ESCC N = 81, PDAC N = 177, UCEC N = 543, BLCA N = 405, BRCA N = 1090, KIRC N = 530, STAD N = 375, CESC N = 304, HNSC N = 500, LIHC N = 371, ESCA N = 80, OV N = 371, LUSC N = 501, LUAD N = 513, SARC N = 259, READ N = 165, THCA N = 502, THYM N = 119.

**Supplemental Figure 7**. **Individual Kaplan-Meier survival curves for M0 macrophage associated ECM signature genes. A-E)** Kaplan-Meier survival curves for **A)** FN1 **B)** COL11A1 **C)** VCAN **D)** MXRA5, **E)** SFRP2, divided by low and high gene expression levels of each individual gene. Serous ovarian cancer patients N = 523. Median survival time is calculated at 50% survival probability.

**Supplemental Figure 8. Overview of the studied ovarian cancer omental samples and their immune landscape. A)** Schematic of ovarian metastatic sample acquisition and analyses. **B)** IHC sections of PAX8 stain increasing with disease score. **C)** Representative digital analysis using Definiens imaging software (Tissue ID: G170). **D)** Ovarian cancer metastatic omentum region of interest (ROI) percentage ordered by increasing disease score (N = 39). **E)** Representative PAX8 stains and matched QuPath digital overlay (Tissue ID: G177). **F)** Scatter plot of PAX8 cell count by QuPath analysis versus disease score. **G)** Representative IHC stains for immune cells in low and high disease tissues. **H)** Cleveland plots of IHC staining for CD68 (N = 38), CD163 (N = 38), CD20 (N = 38), CD3 (N = 38), CD4 (N = 38), CD8 (N = 38) and FOXP3 (N = 38) (cells/ tissue).

**Supplemental Figure 9.** **H&E stains of cellularized and decellularized tissues** from **A-B)** low, medium and high diseased ovarian cancer omental samples. Scale bar: 100 µm.

**Supplemental Figure 10.** **Representative sections of IHC stained low (G159) and high diseased (G273) cellularized and decellularized ovarian cancer samples.** **A)** Stains included anti-VCAN (polyclonal, Ab202906; 1:200), anti-COL1A1 (polyclonal, HPA011795; 1:500), anti-FN1 (polyclonal, Ab23750; 1:500), anti-CTSB (ab125067; 1:50), from Abcam, anti-CS (clone CS-56, Ab11570; 1:600). **B)** Scanning Electron Microscopy (SEM) images of matched cellularized and decellularized omentum samples. Scale bars are indicated on each image. **C)** Representative pseudocolor plots of macrophages recovered from decellularized tissue cultures on day 14. Low disease: G145; high disease: G343.

**Supplemental Figure 11. Optimisation of macrophage cell recovery for flow cytometry analysis.** **A)** Live/ dead analysis of macrophages cultured on general tissue culture plastic (TCP) and in the decellularized tissue model, and the four detachment methods used to recover cells (Cell dissociation buffer, PBS supplemented with 5mM EDTA, PBS supplemented with 0.5%/ 0.2% Trypsin/ EDTA, and Accutase buffer). **B)** Histogram overlays comparing marker expression of macrophages between collection methods and between MO and the decellularized tissue model cultures. Red – trypsin, Green – accutase, Orange – PBS EDTA, and Blue – cell dissociation buffer. N = 2.

**Supplemental Figure 12.** **39 Ovarian cancer (OvCa) patients were used to build the omentum tissue library**. HGSOC: high grade serous ovarian cancer, HGCCC: high grade clear cell cancer, MBT: malignant Brenner tumor.

**Supplemental Figure 13. Overview of the immune landscape. A)** Barplot of mean number of cells/tissue for IHC staining for CD68, CD20, FOXP3, CD4 and CD8 immune cells across all ovarian cancer tissues. N = 38. **B)** Barplot of number of cells/tissue for IHC staining for CD68, CD20, FOXP3, CD4 and CD8 immune cells for each individual ovarian cancer tissue ranked by disease score.

**Supplemental Figure 14. Matrisome heterogeneity of ovarian cancer omentum**. **A)** Protein numbers and total abundances of matrisome categories summed from proteomic enrichment values. Plotted values corresponding to Table A as percentage. N = 39. **B)** Disease score values between ECGs. **C)** Matrisome categories between ECGs. **D)** Scatter plot of PAX8^+^ cells against CD163^+^ cells across OvCa samples. Geometric mean of PAX8^+^ and CD163^+^ values were calculated and indicated by dashed line and defined samples by low or high macrophage counts in context to total samples. **E)** OvCa high disease only samples. Geometric mean PAX8^+^ and CD163^+^ values were calculated and indicated by dashed line in context to high macrophage samples only (from Supplemental Figure 13F). **F)** Scatter plot of PAX8^+^ cells against CD8^+^ cells across OvCa samples. Geometric mean of PAX8^+^ and CD163^+^ values were calculated and indicated by dashed line and defined samples by low or high macrophage counts in context to total samples. **G)** Scatter plot of PAX8^+^ cells against CD4^+^ cells across OvCa samples. Geometric mean of PAX8^+^ and CD163^+^ values were calculated and indicated by dashed line and defined samples by low or high macrophage counts in context to total samples.

**Supplemental Figure 15. Versican and fibronectin are heterogeneous matrisome proteins which associate with increased stroma. A)** Versican and **B)** fibronectin mass ratio values (enrichment) associated with percentage (%) of adipose, stroma and tumour as defined by Definiens® digital analysis. n=38.

**Supplemental Figure 16.** Matrisome molecules associated with M0 macrophages are enriched in cell-derived matrices (CDMs) from patient-derived omental fibroblasts compared to HGSOC metastatic malignant cells. **A)** Schematic summarising the culture methods involved in the generation of CDMs **B)** Table of *in vitro* cultured patient-derived cell types **C)** Malignant and fibroblast CDMs classified into matrisome categories **D)** Volcano plot of p-value and log2FC for matrisome proteins. Red = non-significant with log2 fold change <1 or >-1(NS), blue = non-significant with log2 fold change >1 or <-1 (FC), green = significant with log2 fold change >1 or <-1 (FC_P). **E)** Averaged mass ratio for matrisome protein expression associated with M0 macrophages in malignant cells and fibroblasts. ** p-value < 0.01

**Supplemental Figure 17. Tumor ECM alters the macrophage transcriptome**. **A)** List of decellularized tissues used for RNAseq macrophage cultures. N = 8. **B)** PCA using all genes. **C)** Exploratory boxplot for sample quality, unsupervised cluster analysis and PCA revealed G198_D3 poor quality. **D)** Table of number of differentially expressed (DE) genes identified using the lm model of the limma R package and voom normalization with a ~ECM.type+donor model design. LogFC > 0 denotes up in tumor, logFC < 0 denotes down in tumor. More stringent analysis used fold change greater than 2 (logFC ≥ 1 or logFC ≤ -1). **E)** Volcano plot of p-values and Log_2_FC of tumor verse adjacent tissue cultured macrophages

**Supplemental Figure 18. Tumor extracellular matrix-educated macrophages share signature with M0 macrophages. A)** Stacked bar chart of immune cell predictions by CIBERSORTx grouped by monocytes cultured on tumor or adjacent decellularized tissue. Cell color codes are presented in a key below with color matching across methods. **B)** Barplot of all macrophage subsets and monocytes identified in tumor and adjacent MAM populations from CIBERSORTx analysis on DE genes. Two-way ANOVA with Sidak’s multiple comparison test. ****p < 0.0001.

**Supplemental Figure 19.** **Tumor and adjacent MAMs share similar features with macrophages activated by fatty acid stimuli and LPS stimuli, respectively.** **A)** Heatmap showing correlation between WCGNA modules and genes significantly up-regulated in tumor MAMs compared to adjacent MAMs. Each module of the 49 modules has been ascribed a unique colour and colours name. Red indicates a positive correlation, blue indicates a negative correlation. Significant p-values are indicated. **B)** Correlation of top 4 tumor MAM and adjacent MAM WCGNA modules with biological stimuli.

**Supplemental Figure 20. Day 0 healthy human PBMC derived monocytes used for decellularized tissue culture**. PBMC derived monocytes isolated at day 0 from healthy human blood donors were analyzed by flow cytometry for transmembrane marker expression. Representative histograms of day 0 monocytes marker expression. Same donor monocytes were seeded onto decellularized tissues and later collected and analyzed by RNAseq after 14 days of culture (Figure 4-5).

**Supplemental Figure 21. Tumor ECM alters the macrophage secreted chemokines**. LEGENDplex™ quantification (pg/mL) of chemokine levels in macrophage supernatants from tumor and adjacent ECMcultured macrophages. N = 8. Duplicates used for each donor.

Supplemental Video 1. **Representative video of laser capture dissection.** The video shows tumor areas being cut and selected on a PALM dissection microscope,
