## Supplemental Figures for "Extracellular matrix educates a tumor macrophage phenotype found in ovarian cancer metastasis"

Supplemental Figure 1.

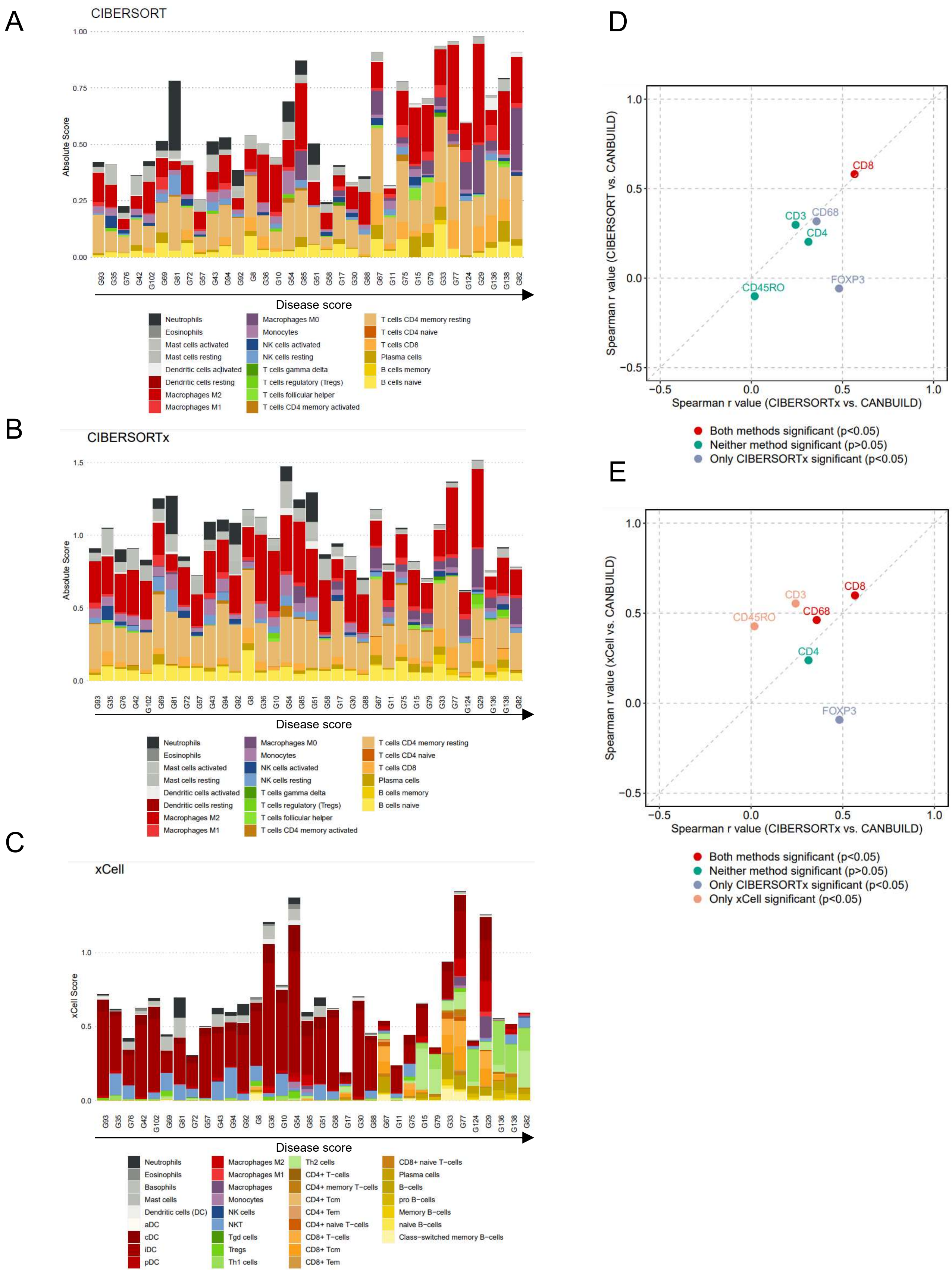

### Supplemental Figure 2.

A

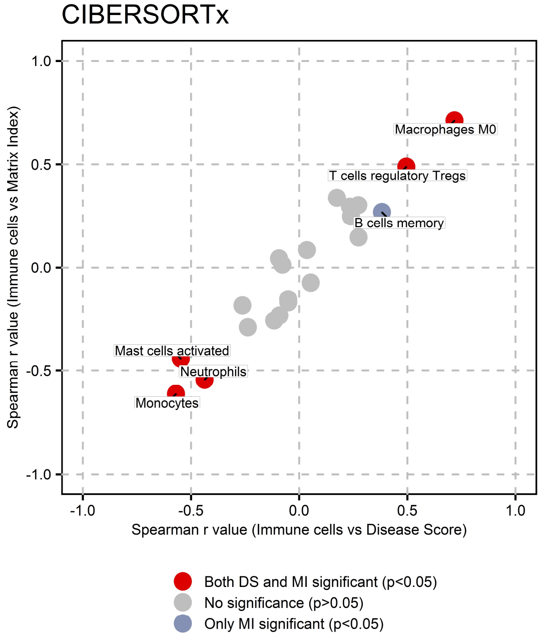


B

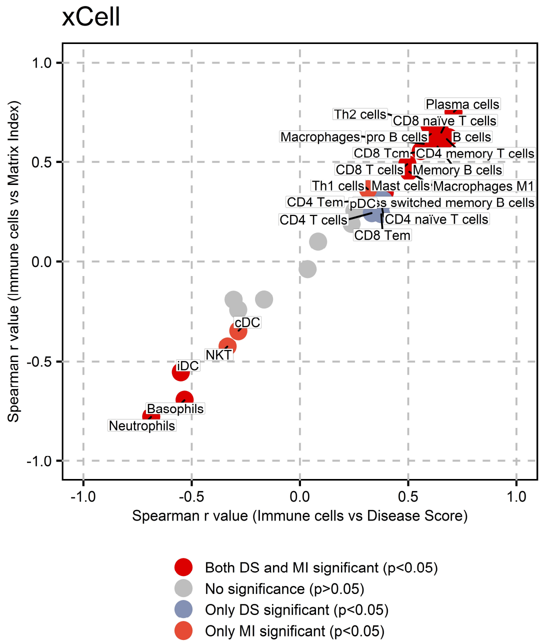

Supplemental Figure 3.

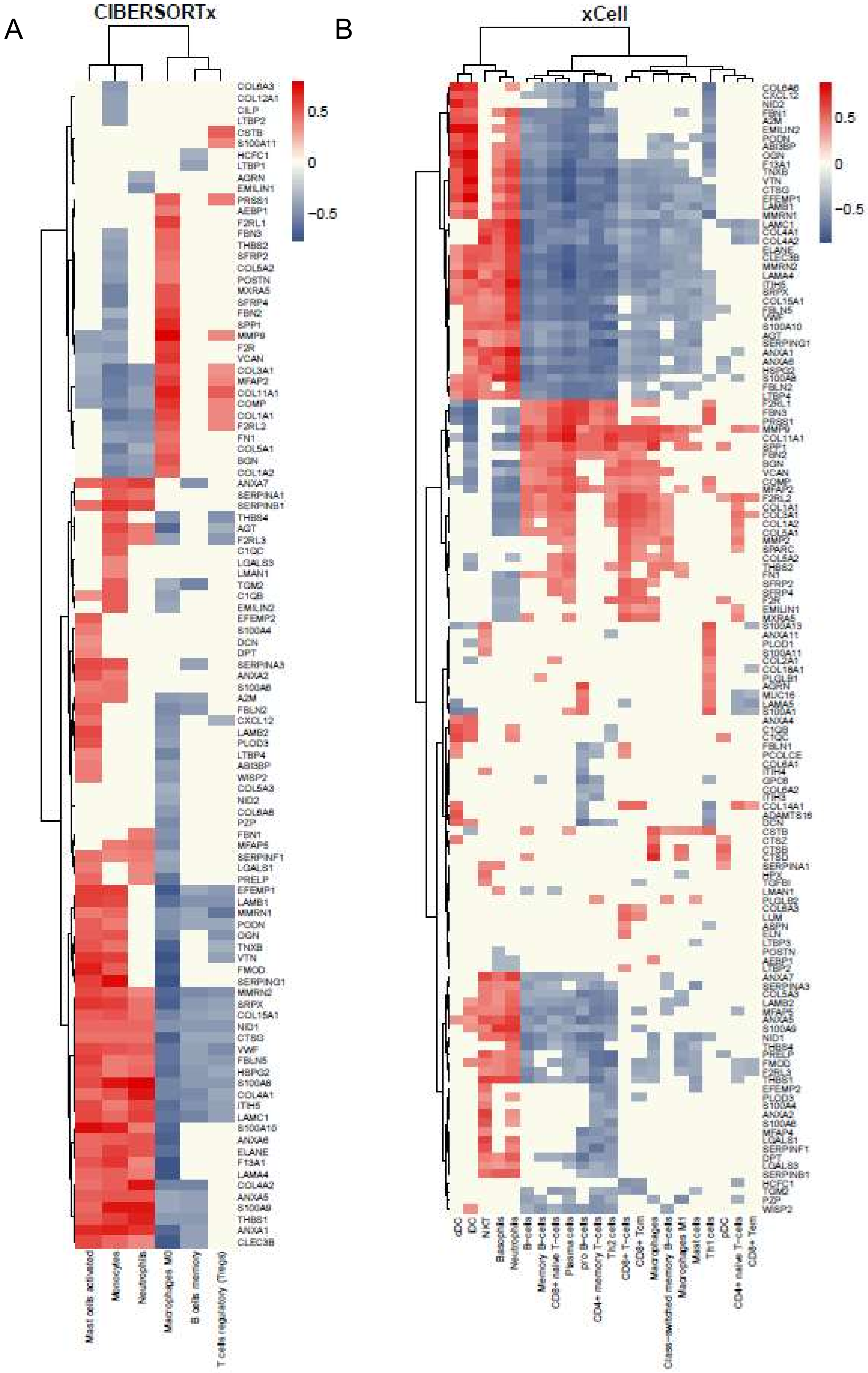

Supplemental Figure 4.

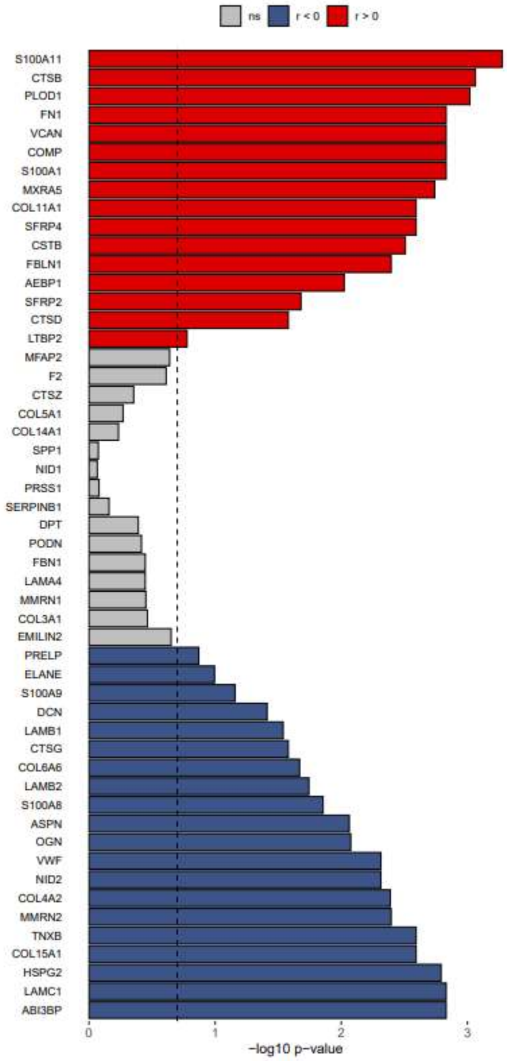

Supplemental Figure 5.

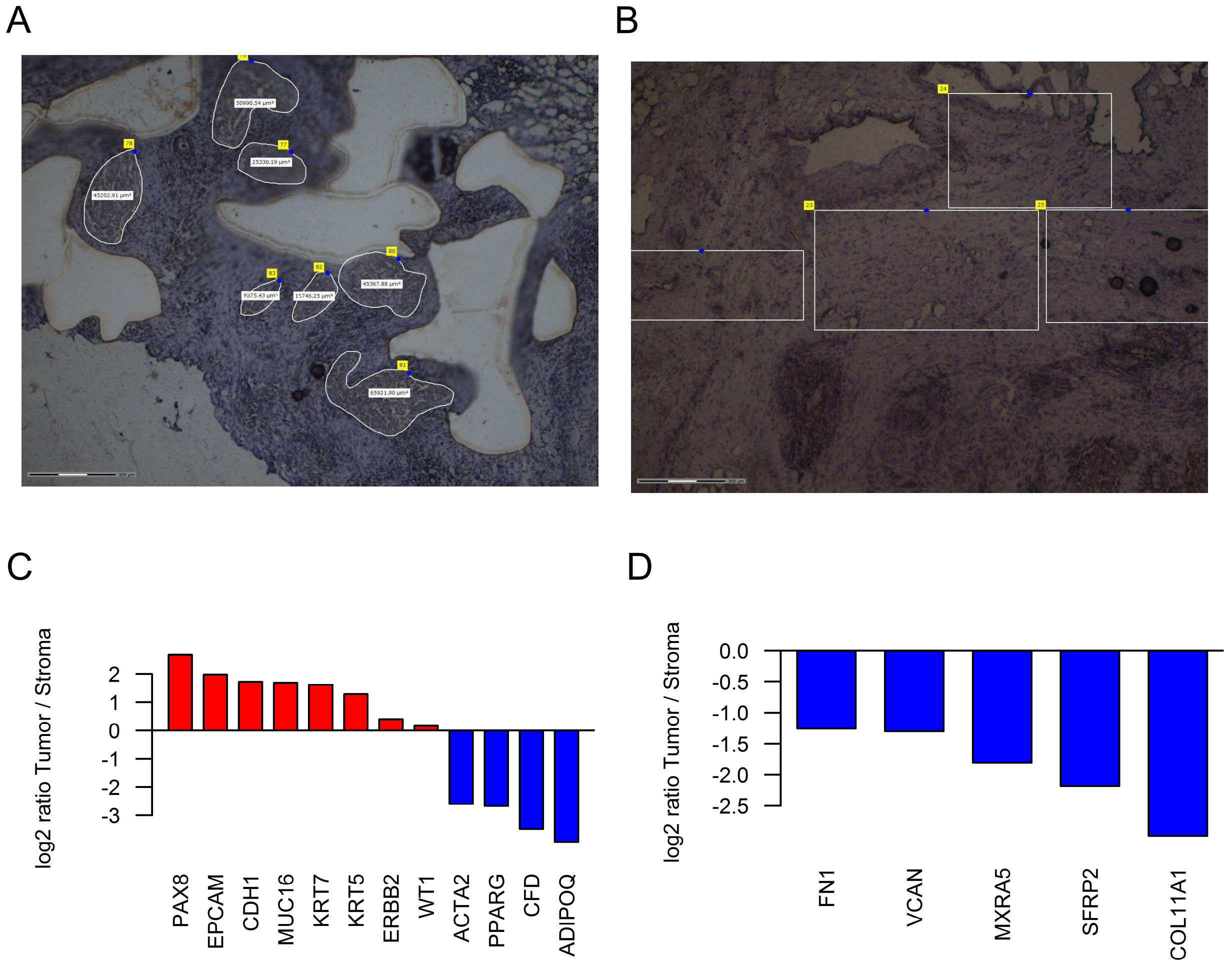

### Supplemental Figure 6.

A

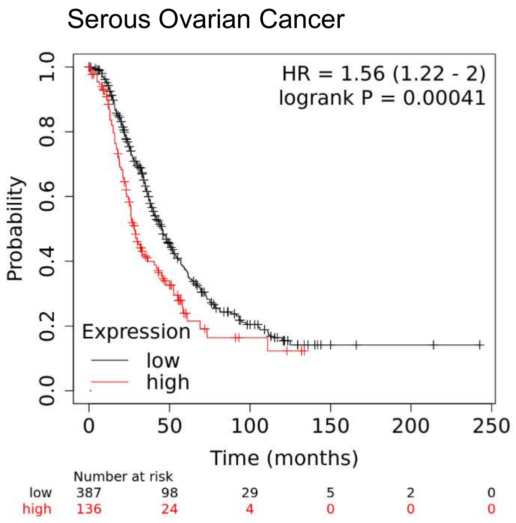


Median Overall Survival

| Low expression cohort (months) | High expression cohort (months) |
|---|---|
| 45.63 | 28.25 |

B

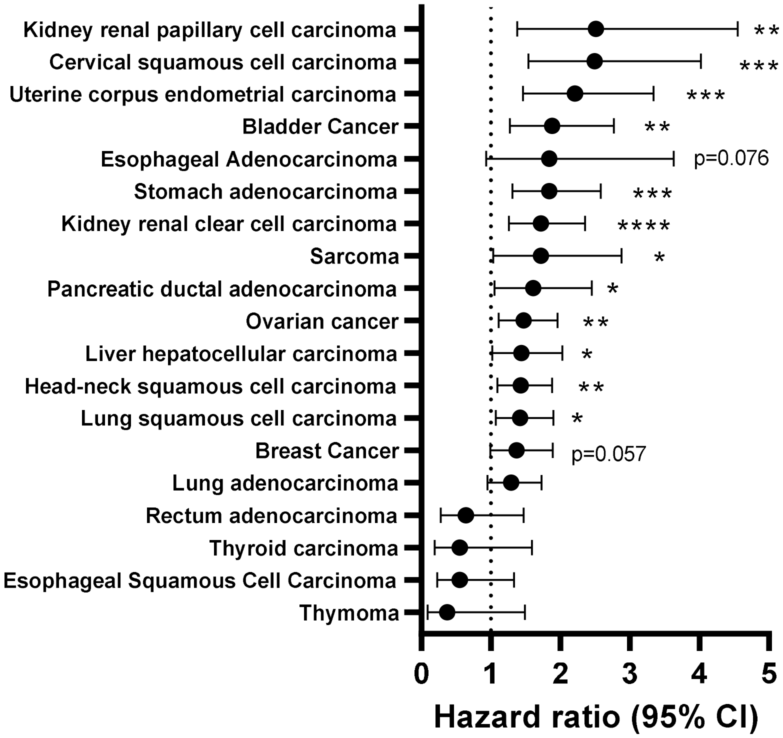

Supplemental Figure 7.

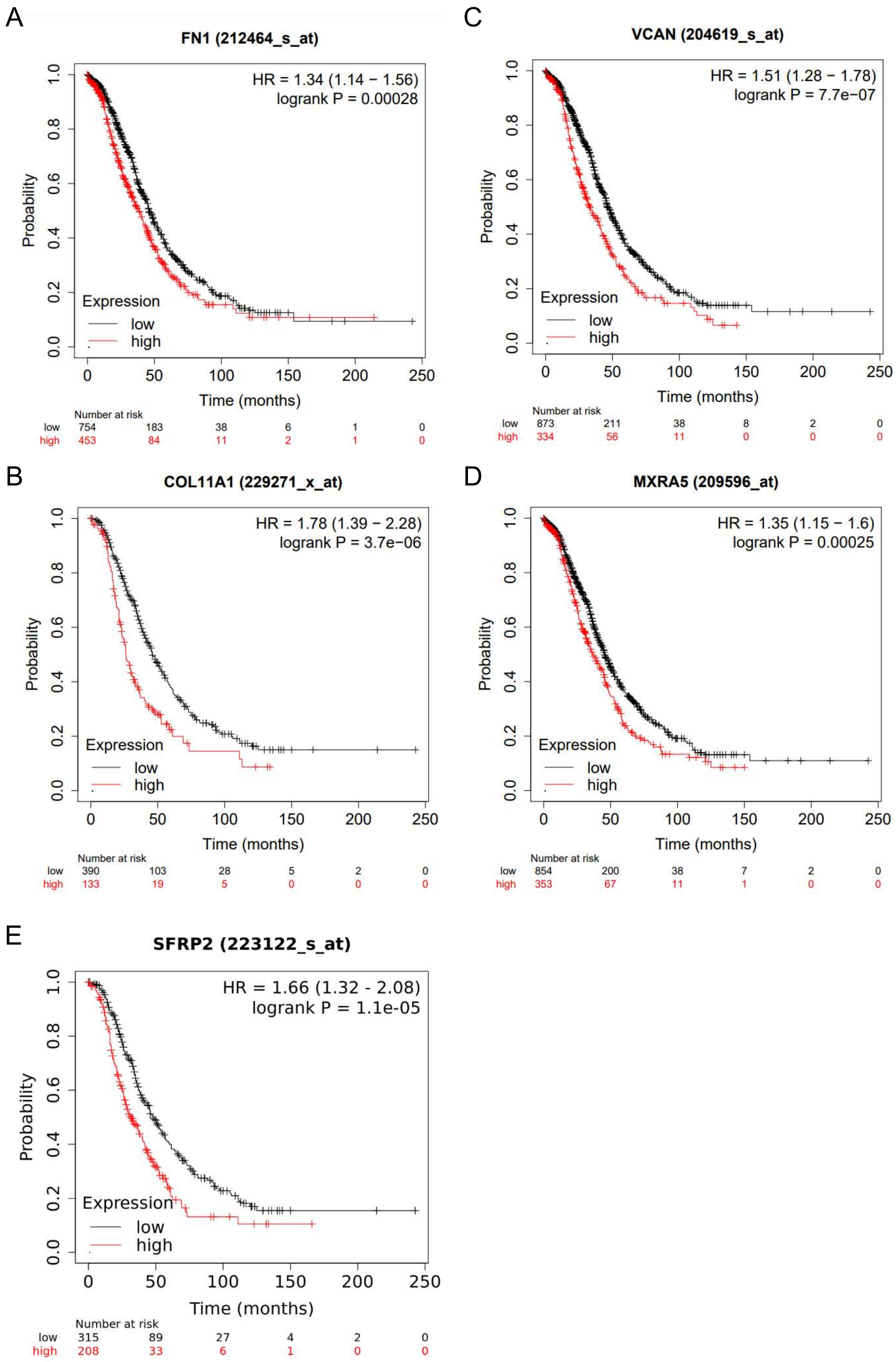

Supplemental figure 8.

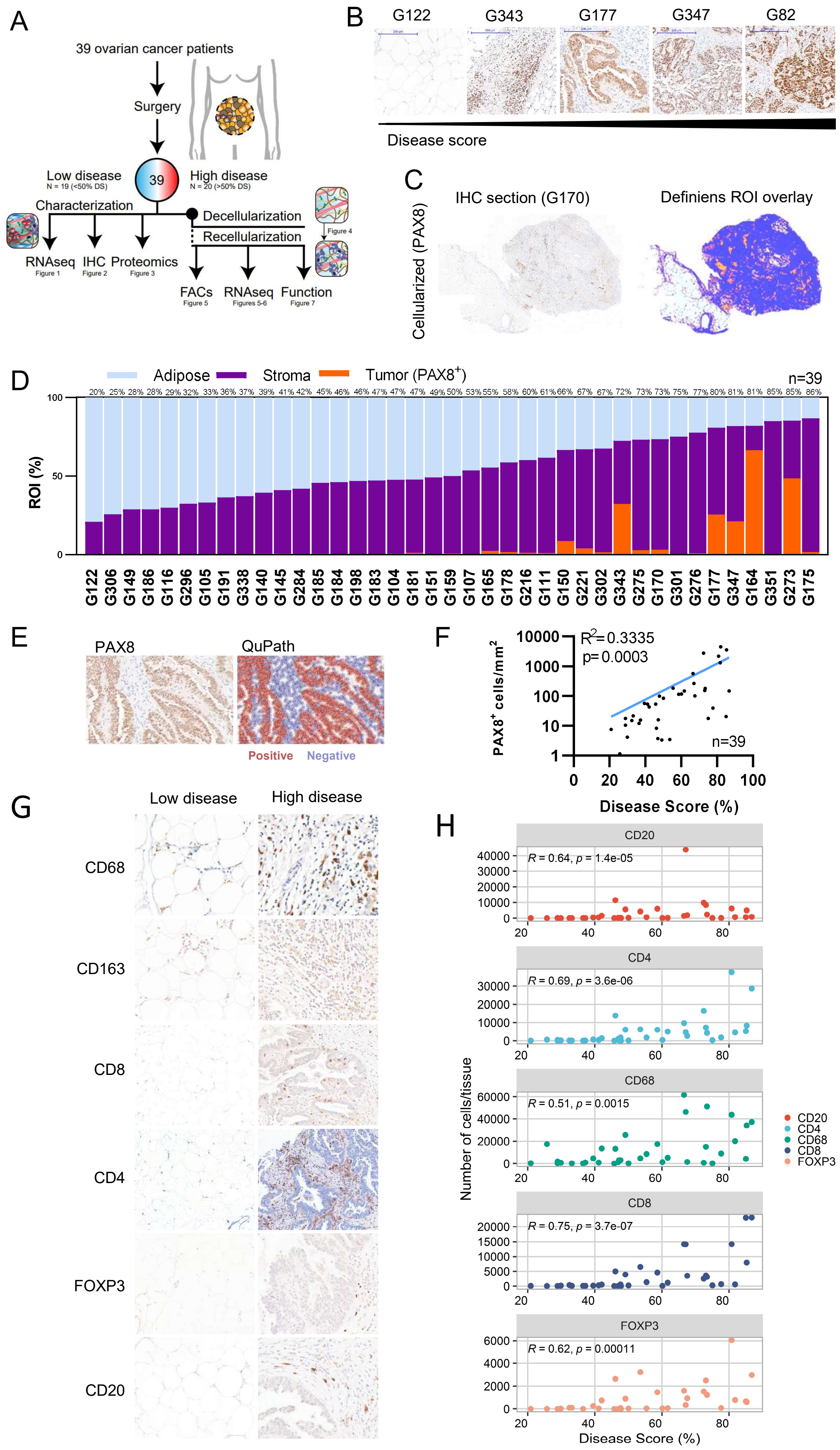

Supplemental Figure 9.

A

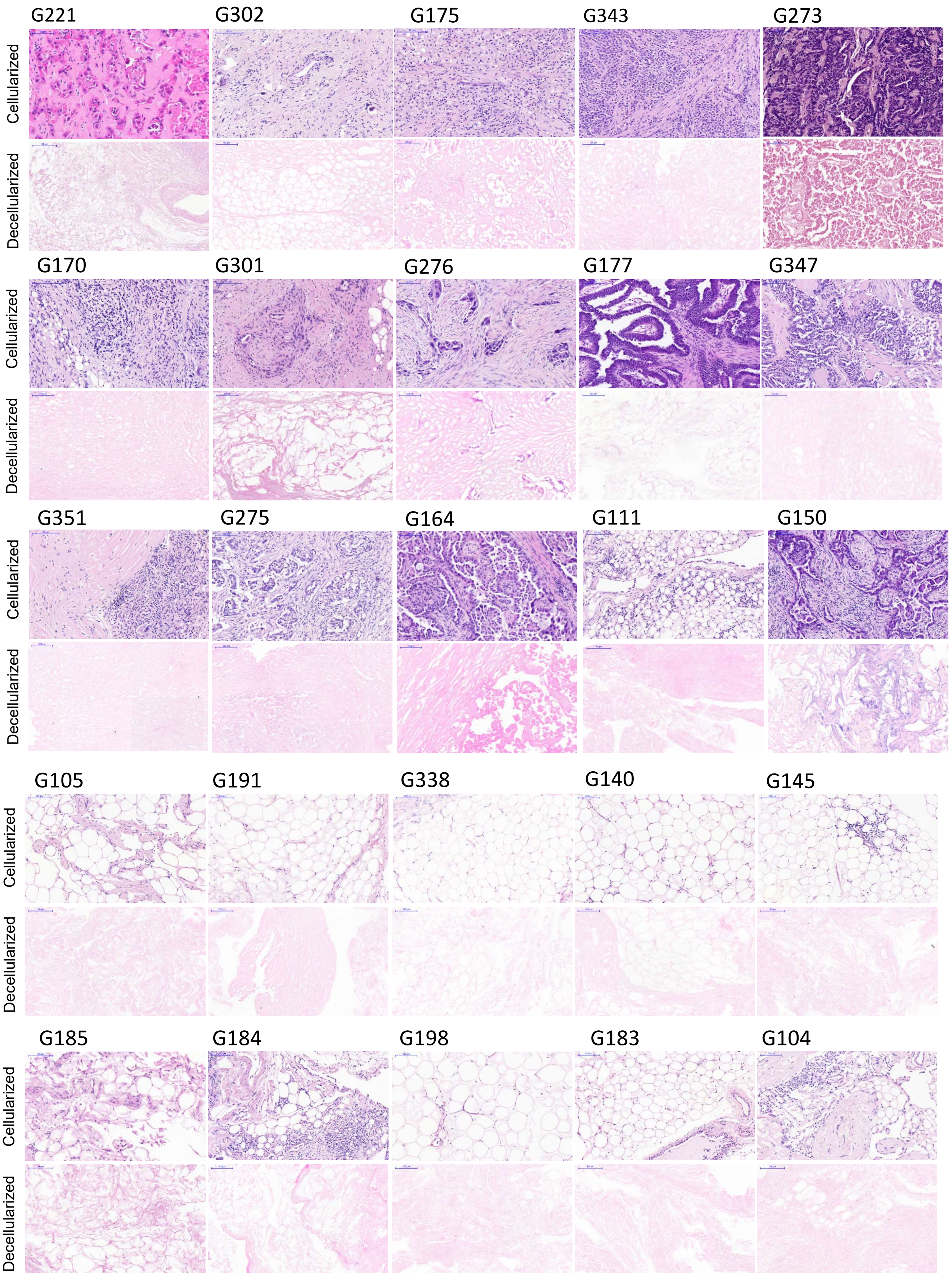

Supplemental Figure 9.

B

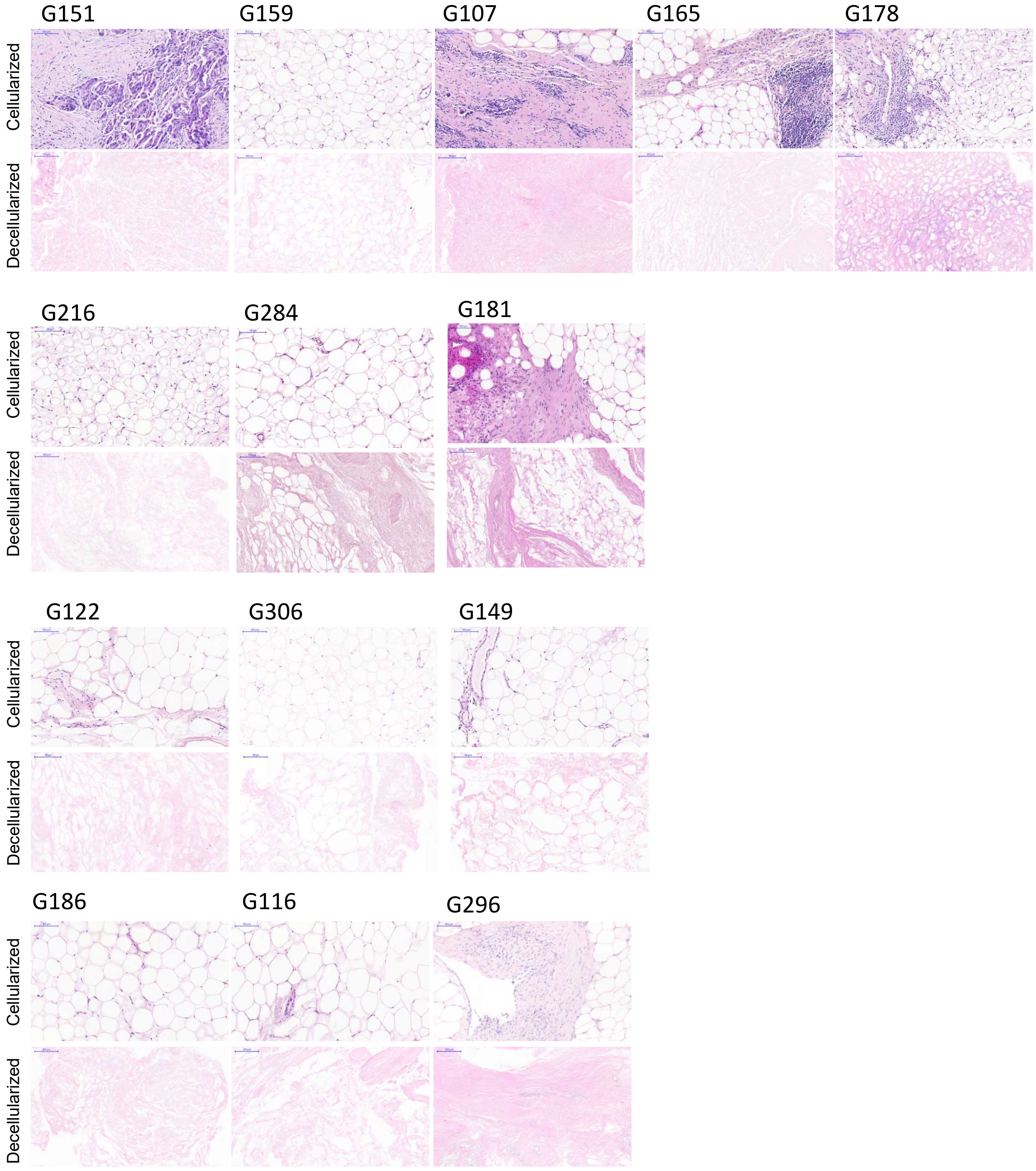

Supplemental Figure 10.

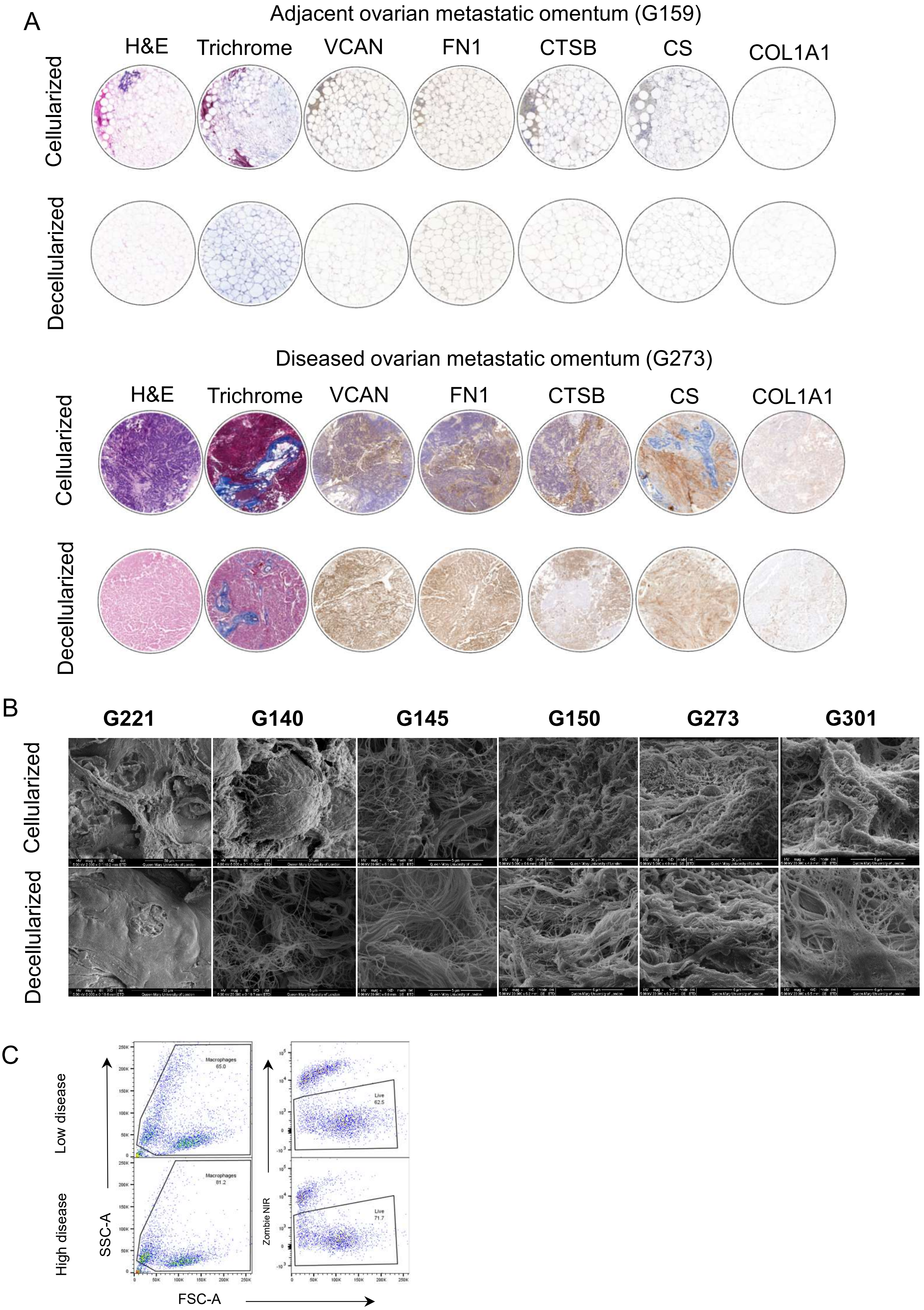

Supplemental Figure 11.

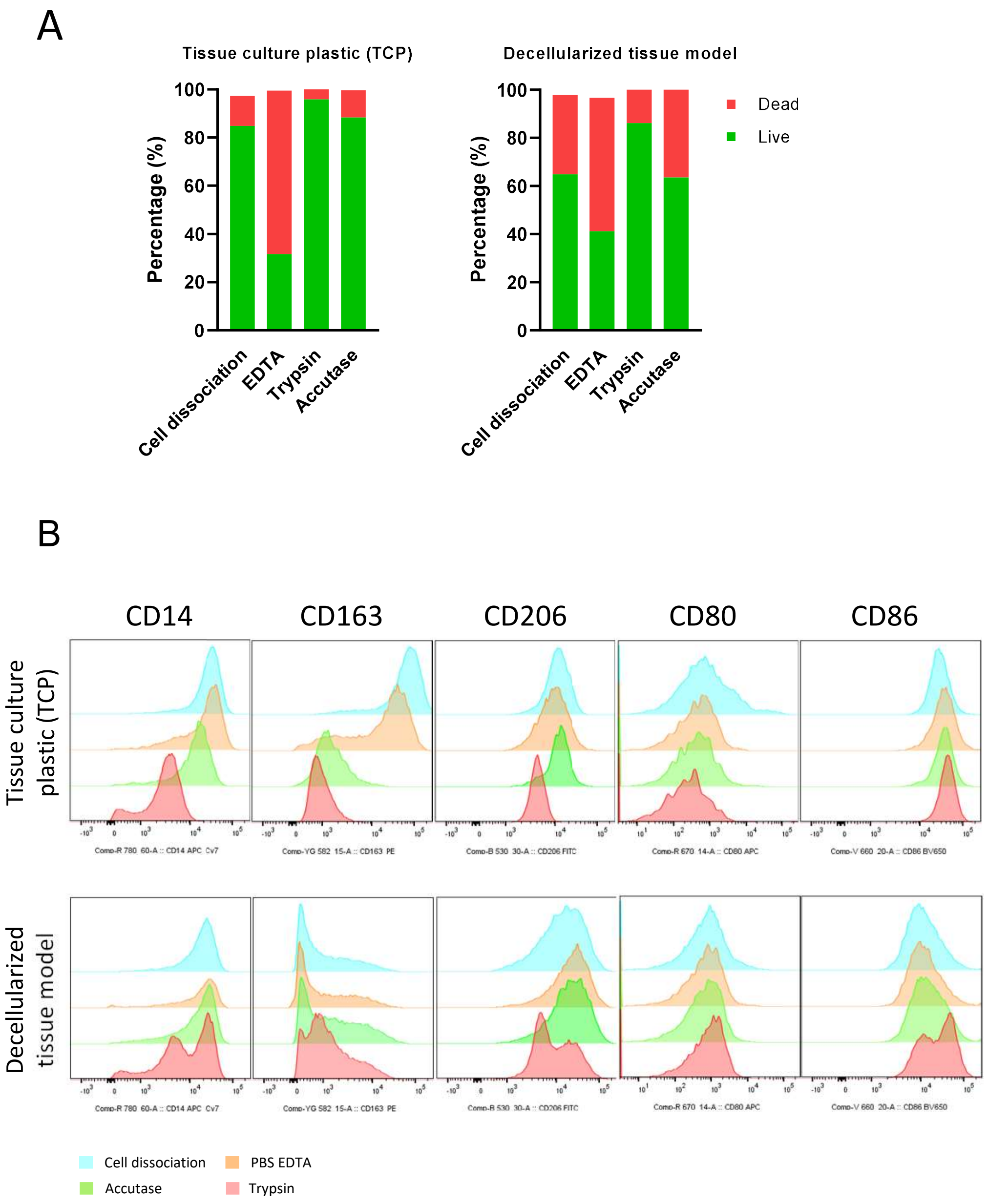

Supplemental Figure 12.

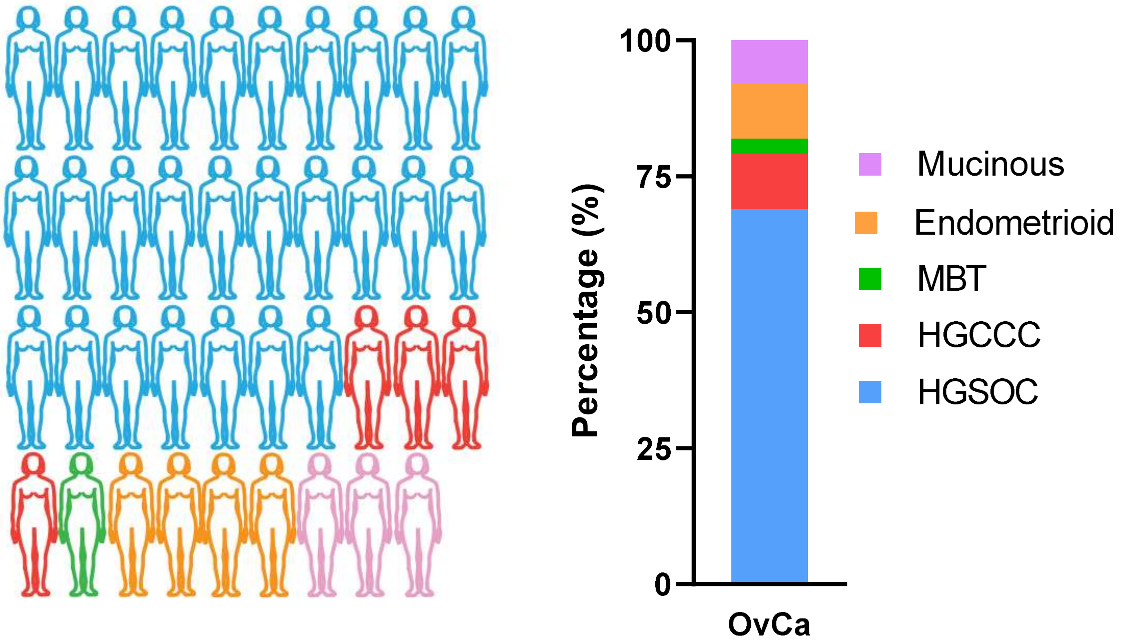

### Supplemental Figure 13.

A

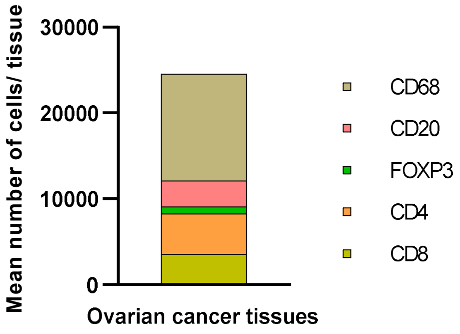


B

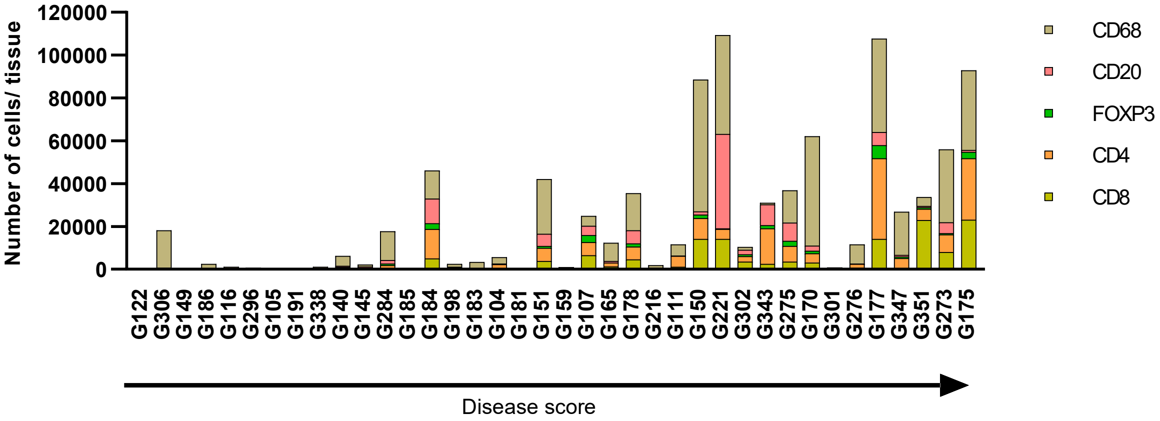

### Supplemental Figure 14.

A

| | Protein numbers | Total abundance | Total abundance (%) |
|---|---|---|---|
| Collagens | 29 | 4.E+06 | 35.49 |
| Glycoproteins | 96 | 3.E+06 | 25.25 |
| Regulators | 95 | 9.E+05 | 7.93 |
| Affiliated | 52 | 6.E+05 | 4.89 |
| Proteoglycans | 16 | 3.E+06 | 25.29 |
| Secreted factors | 50 | 1.E+05 | 1.12 |

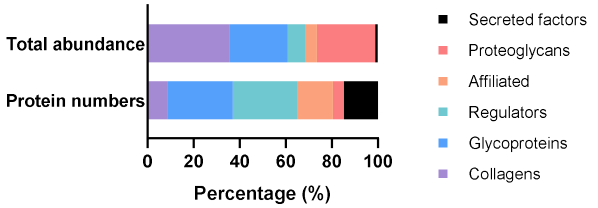


B

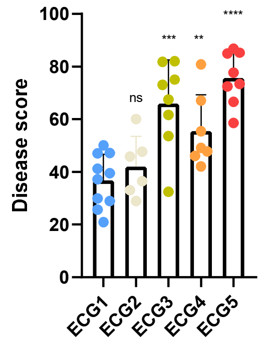


C

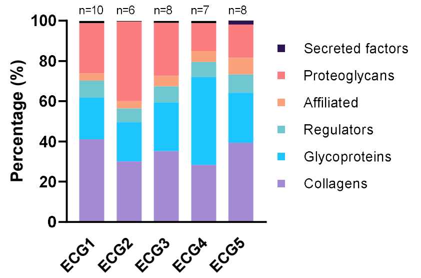


D

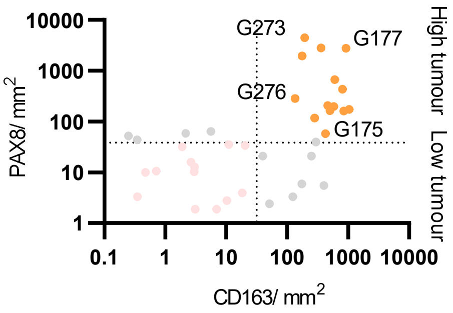


E

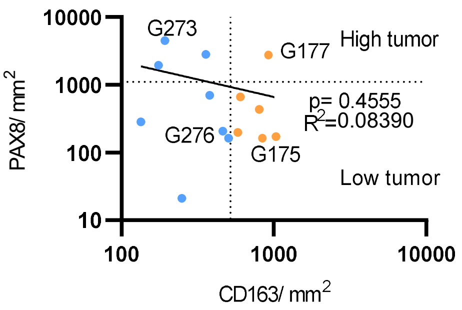


F

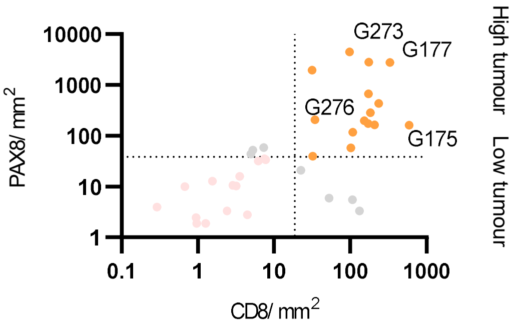


G

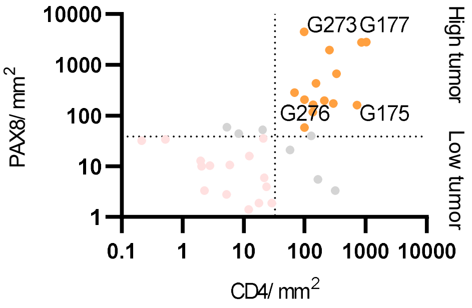

### Supplemental Figure 15.

A

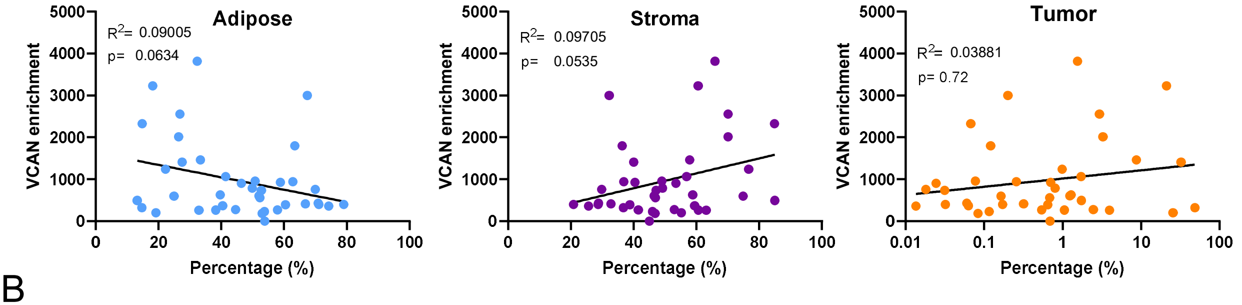


B

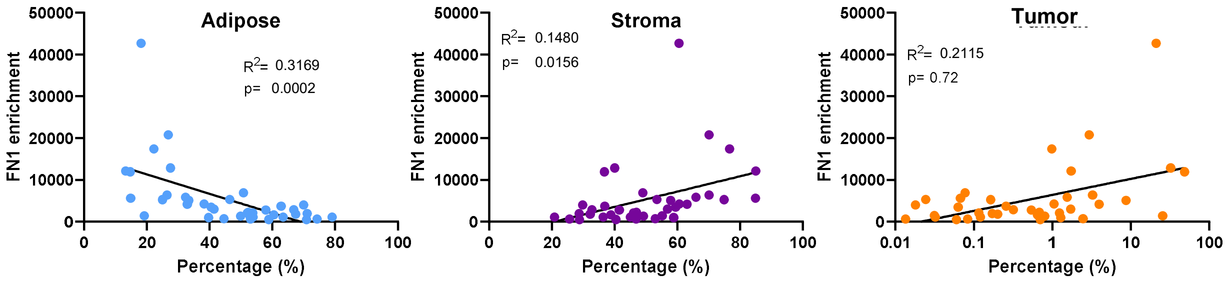

Supplemental Figure 16.

A

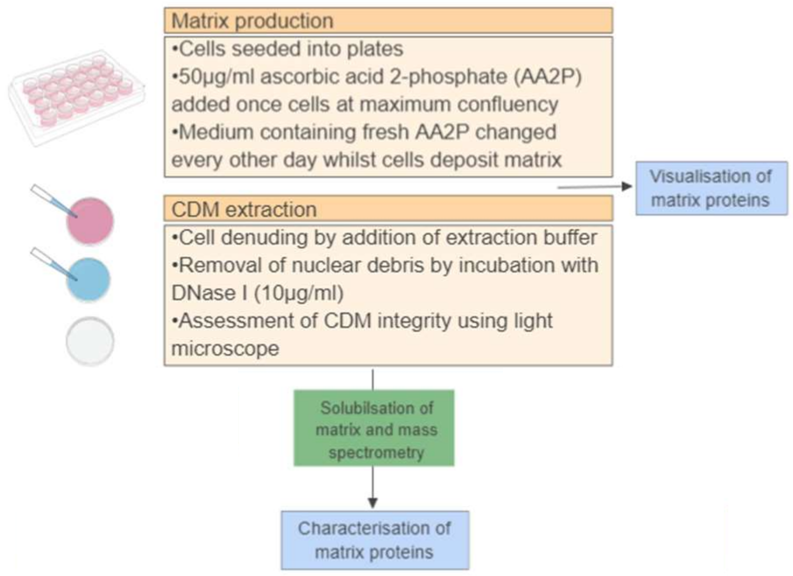


B

| ID | Cell | Type/area |
|---|---|---|
| AOCS1 | Malignant | Metastasis |
| G164 | Malignant | Metastasis |
| G342 | Fibroblast | Omentum |
| G351 | Fibroblast | Omentum |
| G369 | Fibroblast | Omentum |

C

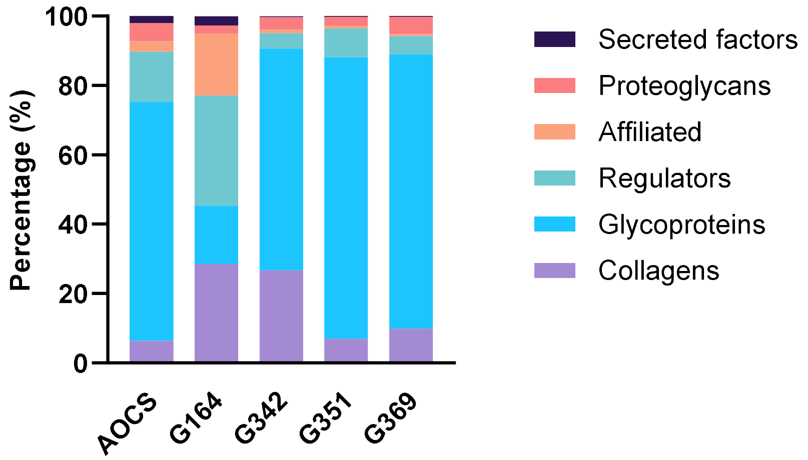


E

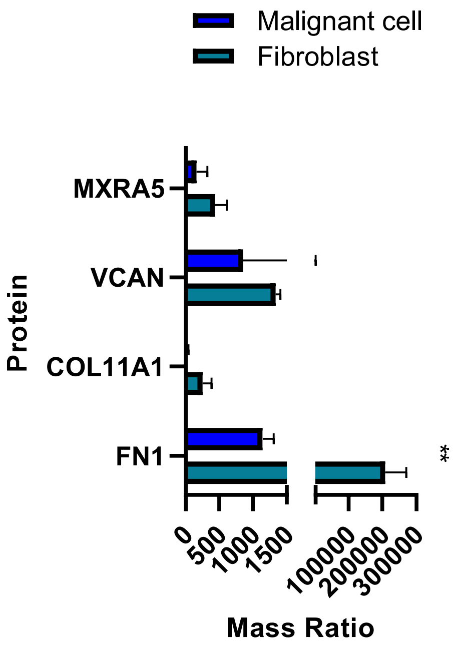


D

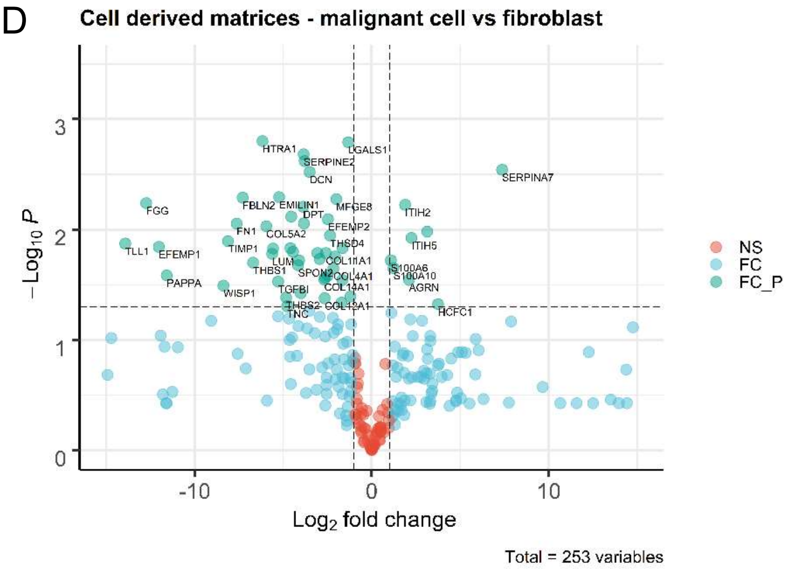

### Supp Figure 17.

A

| ID | Type | Disease score |
|---|---|---|
| **G164** | Tumor | 81.96013 |
| **G302** | Tumor | 67.54765 |
| **G276** | Tumor | 77.65805 |
| **G150** | Tumor | 66.59231 |
| **G122** | Adjacent | 20.91347 |
| **G306** | Adjacent | 25.73339 |
| **G198** | Adjacent | 46.95865 |
| **G145** | Adjacent | 41.16155 |

B

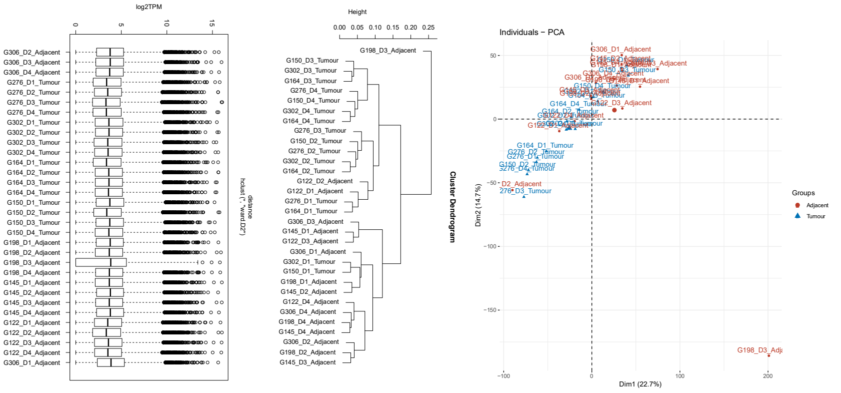


C

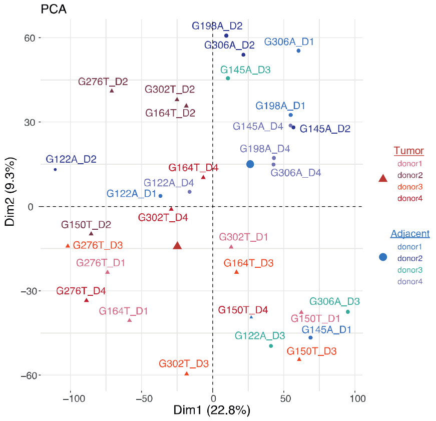


D

| Thresholds | Number of DE genes |
|---|---|
| Protein-coding & adj. p. val =< 0.05 & logFC>=0 | 1839 |
| Protein-coding & adj. p. val =< 0.05 & logFC=<0 | 1774 |
| Protein-coding & adj. p. val =< 0.05 & logFC>=1 | 261 |
| Protein-coding & adj. p. val =< 0.05 & logFC>=-1 | 198 |

E

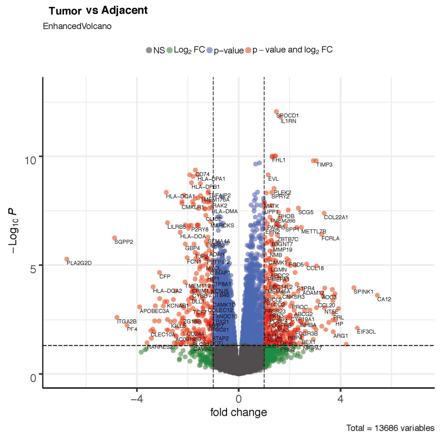

### Supplemental Figure 18

A

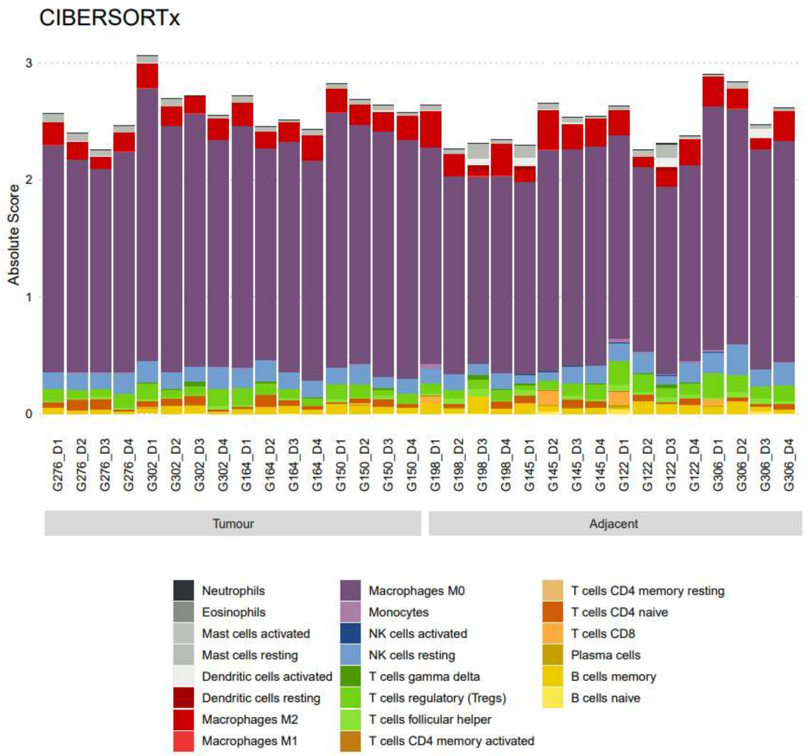


B

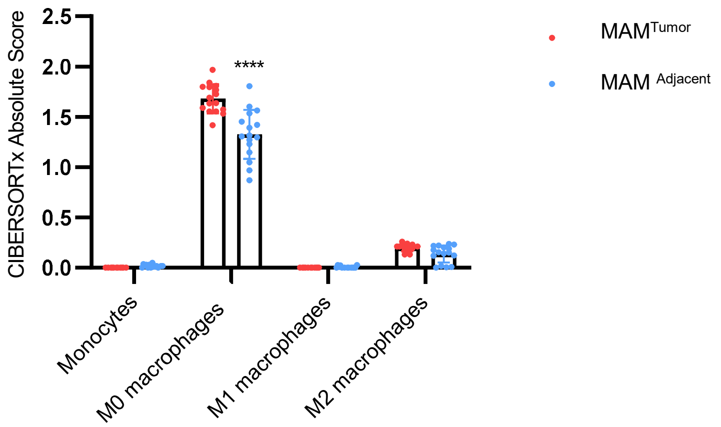

### Supplemental Figure 19

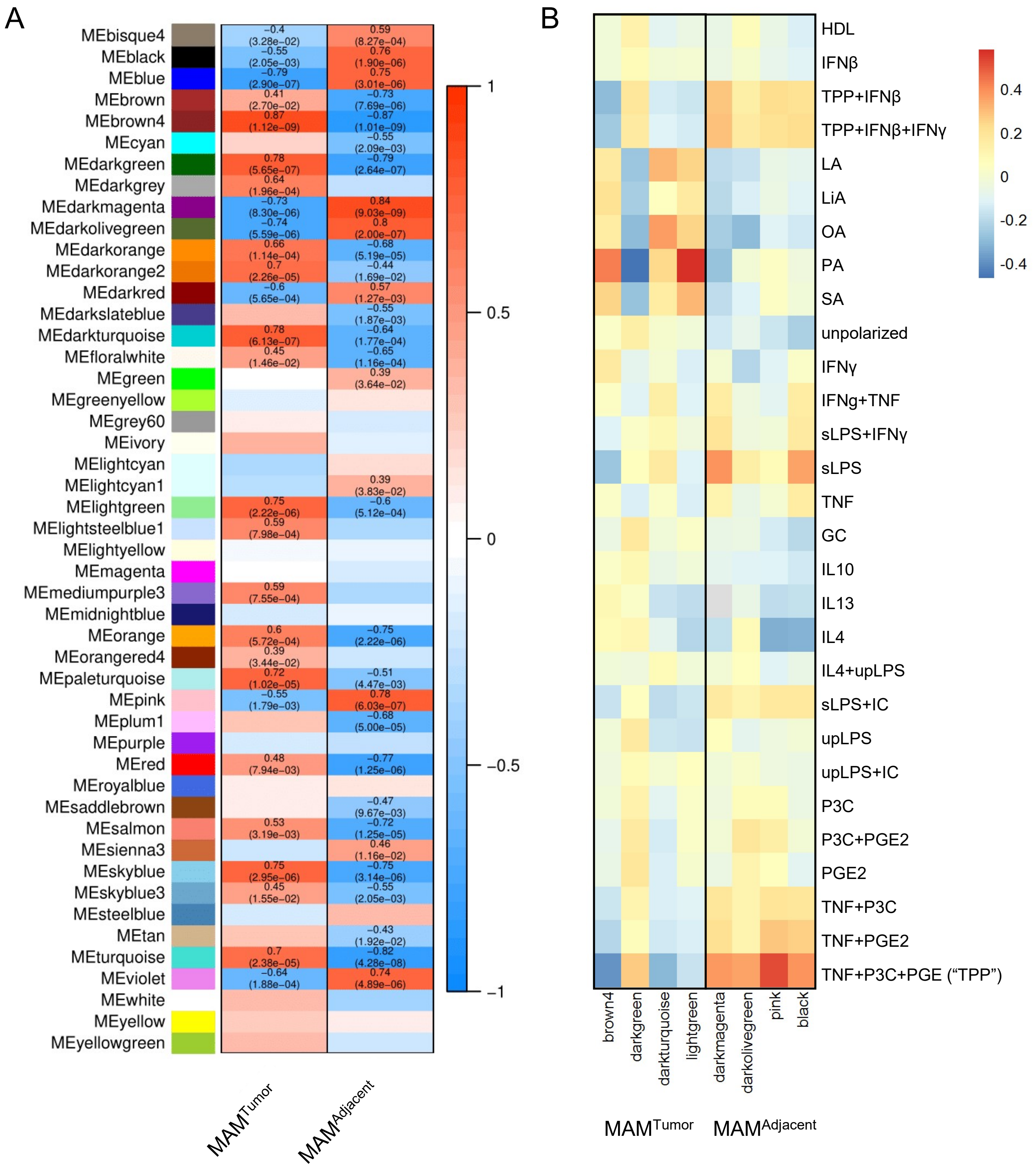

### Supp Figure 20.

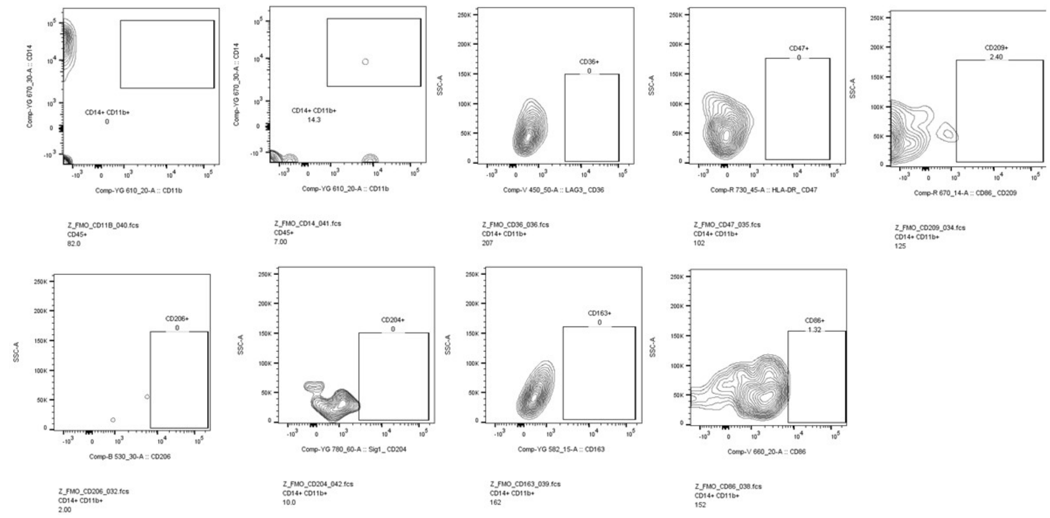

### Supp Figure 21.

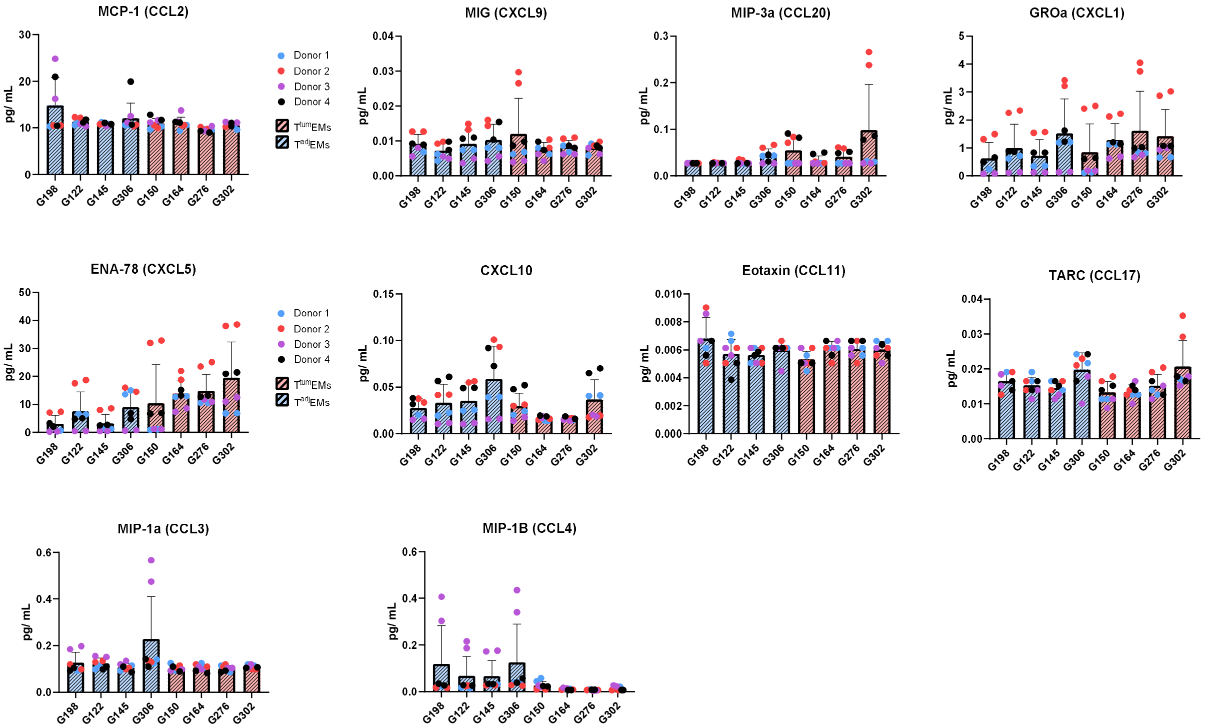
